# Evolutionary dynamics in structured populations under strong population genetic forces

**DOI:** 10.1101/579854

**Authors:** Alison F. Feder, Pleuni S. Pennings, Joachim Hermisson, Dmitri A. Petrov

## Abstract

High rates of migration between subpopulations result in little population differentiation in the long-term neutral equilibrium. However, in the short-term, even very abundant migration may not be enough for subpopulations to equilibrate immediately. In this study, we investigate dynamical patterns of short-term population differentiation in adapting populations via stochastic and analytical modeling through time. We characterize a regime in which selection and migration interact to create non-monotonic patterns of the population differentiation statistic *F_ST_* when migration is weaker than selection, but stronger than drift. We demonstrate how these patterns can be leveraged to estimate high migration rates that would lead to panmixia in the long term equilibrium using an approximate Bayesian computation approach. We apply this approach to estimate fast migration in a rapidly adapting intra-host Simian-HIV population sampled from different anatomical locations. Notably, we find differences in estimated migration rates between different compartments, all above *N_e_m* = 1. This work demonstrates how studying demographic processes on the timescale of selective sweeps illuminates processes too fast to leave signatures on neutral timescales.

---

The physical structure of a population can profoundly impact its evolution. Structure can speed up or slow down evolution [1, 2, 3] and cause populations to reach evolutionary outcomes unlikely or impossible in well-mixed populations [4, 5, 6], including the maintenance of genetic diversity [7,8] and the prevalence of cooperation [9], clonal interference [10] and speciation [11]. Understanding population structure aids our interpretation of genetic data sampled from around the globe [12,13,14]. For these reasons, understanding and quantifying population structure has long been a goal of population biology.

Despite population structure’s importance, classical population genetics results suggest that little migration among subpopulations destroys genetic evidence of its existence. Under neutral evolution, long-term population differentiation is a function of the product of effective population size (*N_e_*) and migration rate per individual in the population (*m*). Specifically, Wright seminally showed that population differentiation measure *F_ST_* = 1/(1 + 4*N_e_m*) [15] in an island model under neutral equilibrium. Therefore we might expect that when *N_e_m* is much smaller than one, local drift overwhelms migration and *F_ST_* is elevated above zero (with up to a maximum value of one under certain conditions) and populations appear spatially structured. However, when *N_e_m* is much larger than one, migration is effectively much faster than drift, *F_ST_* approaches zero and the populations appear “effectively panmictic.” Although the island model framework has been criticized [16], the result is a population genetics classic and underlies all population genetics theory (and subsequent applications to data) under the modeling assumption of panmixia. Examples include detecting barriers to gene-flow by using patterns of local divergence to deduce locally reduced levels of genetic exchange.

Importantly, Wright’s equation describes population equilibrium. In such an equilibrium, even very few migrants can mix a large population if gene flow occurs over timescales of *N_e_* generations. For example, in a population of *N_e_* = 10^5^, it makes little difference whether subpopulations are connected by 10 migrants per generation or 10^4^ - the subpopulations will appear near equivalently identical. However, a different picture emerges over shorter timescales (i.e., less than N_e_ generations) where ecological processes can have considerable non-equilibrium effects. Imagine such an ecological disturbance in a population that drastically and immediately perturbs allele frequencies in different demes connected by N_e_m migrants per generation. Even though migration rates *N_e_m* = 10 and *N_e_m* = 10^4^ would both lead to long-term panmixia, these rates result in vastly different amounts of time until the subpopulations equilibrate back to panmixia.

Non-instantaneous (but still rapid) perturbations in allele frequency that occur on a timescale much shorter than 1/*N_e_m* are expected to leave transient patterns in population differentiation. Strong positive selection represents one such process. Importantly, this holds not only for heterogeneous selection (which will lead to permanent differentiation patterns), but also for spatially homogeneous selection, which we will focus on here.

All this to say, migration rates that would result in equivalent panmixia between subpopulations over long time periods (i.e., *N_e_m* ≫ 1) may result in more or less genetic differentiation over short timescales. Most straightforwardly, this presents a problem for estimation of *N_e_m* from *F_ST_*. Fast processes lead to signals of isolation which can be misinterpreted as weak migration or local adaptation preventing gene flow if the populations are assumed to be at equilibrium [2]. However, this also presents an opportunity: with dynamical data it may be possible to track non-equilibrium behavior of population differentiation over time after a strong perturbation to perform migration estimation impossible with Wright’s formula because such rates would result in effectively panmictic *F_ST_* over long time scales.

Here, we show that patterns in population differentiation can result even for very large migration rates if the selection pressure is sufficiently strong. We focus on intrapatient viral adaptation to drugs as a motivating example. Here, we know that adaptation can happen on the timescale of tens of generations and therefore very large (i.e., “effectively panmictic”) migration rates between different tissues or organs can potentially generate very different outcomes with respect to adaptive responses in different parts of the body [17]. Similar patterns over slightly longer time scales can be observed in response to immune selection rather than drug pressure [18]. We examine short timescales consistent with the clinical sampling schedules common in patients.

In this paper, we develop intuitions about subdivided populations adapting rapidly to a new selective pressure and connected to each other by migration over short timescales. Although our theory is more general, we start with a concrete example of how Simian-HIV evolves drug resistance across multiple tissues. This example demonstrates how strong selection, abundant mutation and fast migration can interact to create patterns of substantial yet transient population differentiation over short timescales. We investigate the parameter regime where these forces are jointly strong and find that when migration is significantly faster than drift but slower than selection, characteristic, non-monotonic patterns of population differentiation emerge. We develop an approximate Bayesian computation (ABC) approach that uses these patterns of dynamic population differentiation over time to estimate migration rates far above levels possible to determine with drifting alleles over long timescales. Finally, we return to the Simian-HIV data and estimate between tissue migration rates unresolvable in the neutral equilibrium regime.

## Results

### 1. A motivating example using Simian-HIV drug resistance evolution

Viruses infecting a multicellular host may be subject to both strong selection (via drugs and the immune system) and also fast migration between anatomical compartments (via the circulatory and/or lymphatic systems).

A previous study found that among Simian-HIV within drug-treated macaques, multiple drug resistance mutations spread simultaneously across anatomical compartments with weak but significant evidence of population structure dynamically changing over time [17]. In Figure 1, we reproduce a simplified picture of this evolutionary process in one Simian-HIV-infected macaque (T98133) across three well-sampled tissues (lymph node, gut and plasma) at four time points (1, 3, 8 and 14 weeks after the onset of selection via the drug FTC). Approximately 30 single-genome sequences of the reverse transcriptase region of the *pol* gene are taken per time point, per sampling location.

**Figure 1.**
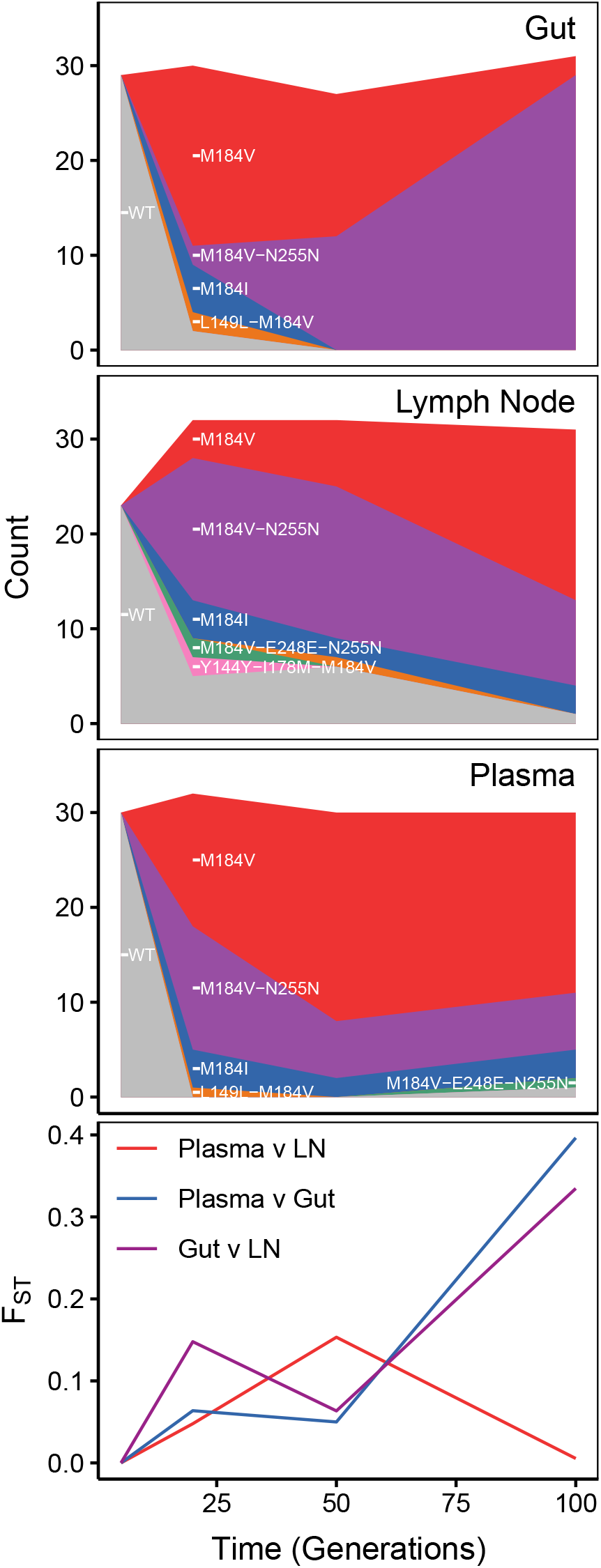
Dynamics of drug resistance fixation across space and time in a treated Simian-HIV population. The top three rows shows diagrams of drug resistant haplotypes spreading in different sampling locations over time sampled at generations 7, 21,49 and 98 after the onset of selection via the drug FTC in the gut, lymph node and blood plasma. Each color represents a distinct lineage separated by at least one mutation. The bottom-most panel plots pairwise *F_ST_* between pairs of sampling locations.

Each population has several different variants of drug resistant viruses spreading simultaneously which can be seen both through different encodings of drug resistant types (i.e., M184V versus M184I) and also through linkage to different hitchhiking mutations (M184V versus M184V + N255N + D177N) (Figure 1). Multiple spreading mutations suggest that the population mutation rate (*θ*) is sufficiently large so as to produce soft selective sweeps [19, 20]. These mutations quickly displace wildtype virus across tissues almost entirely, confirming that selection to drug resistance is strong. Finally, migration is sufficiently fast to spread the same mutations around to different compartments. While some of these apparently spreading drug resistance mutants may be recurrent mutations, several are linked to multiple putatively neutral mutations, suggesting that they arose once and then moved between anatomical compartments. However, migration is not so fast that each compartment looks equivalent, as demonstrated by statistically significant elevated pairwise *F_ST_* between compartments at some of the points during the experiment (Figure 1D) (see Materials and Methods for details, and [17] for a much more extensive description of population differentiation in this data). Notably, we observe non-monotonic patterns of *F_ST_* over time, suggesting that some pairs of populations differentiate as they fix beneficial mutations, then re-equilibrate over time (plasma v. lymph node and lymph node v. gut).

These patterns suggest qualitatively that the population genetic forces of mutation, migration and selection are all jointly strong. However, it remains quantitatively unclear how strong they are with reference to each other. We therefore explore the dynamics of populations over short periods of time (including non-monotonicity of *F_ST_*) to better understand quantitatively how these population genetic parameters interact. We use many of the sampling attributes from the Simian-HIV example (≈100 generations, sampling depth of ≈30 genomes, ≈4 time points) as our reference in exploring this parameter space.

Although we use this data is a motivating example, many ecologically important processes feature rapidly adapting and dynamically interconnected populations.

### 2. Building intuition about the interaction between strong selection and migration

We next explore how rapid adaptation in two subpopulations connected with migration can lead to signals of population differentiation over time.

Figure 2 shows two schematic scenarios through which a population divided into two subpopulations can fix beneficial variants. In both instances, a beneficial mutation will enter one population and begin to spread. However, the paths differ according to whether the first beneficial allele in the second subpopulation arrives through de novo mutation or via migration. Understanding differences in static signatures resulting from these two scenarios (in addition to selection from shared standing genetic variation between the two populations) are considered in [21].

**Figure 2.**
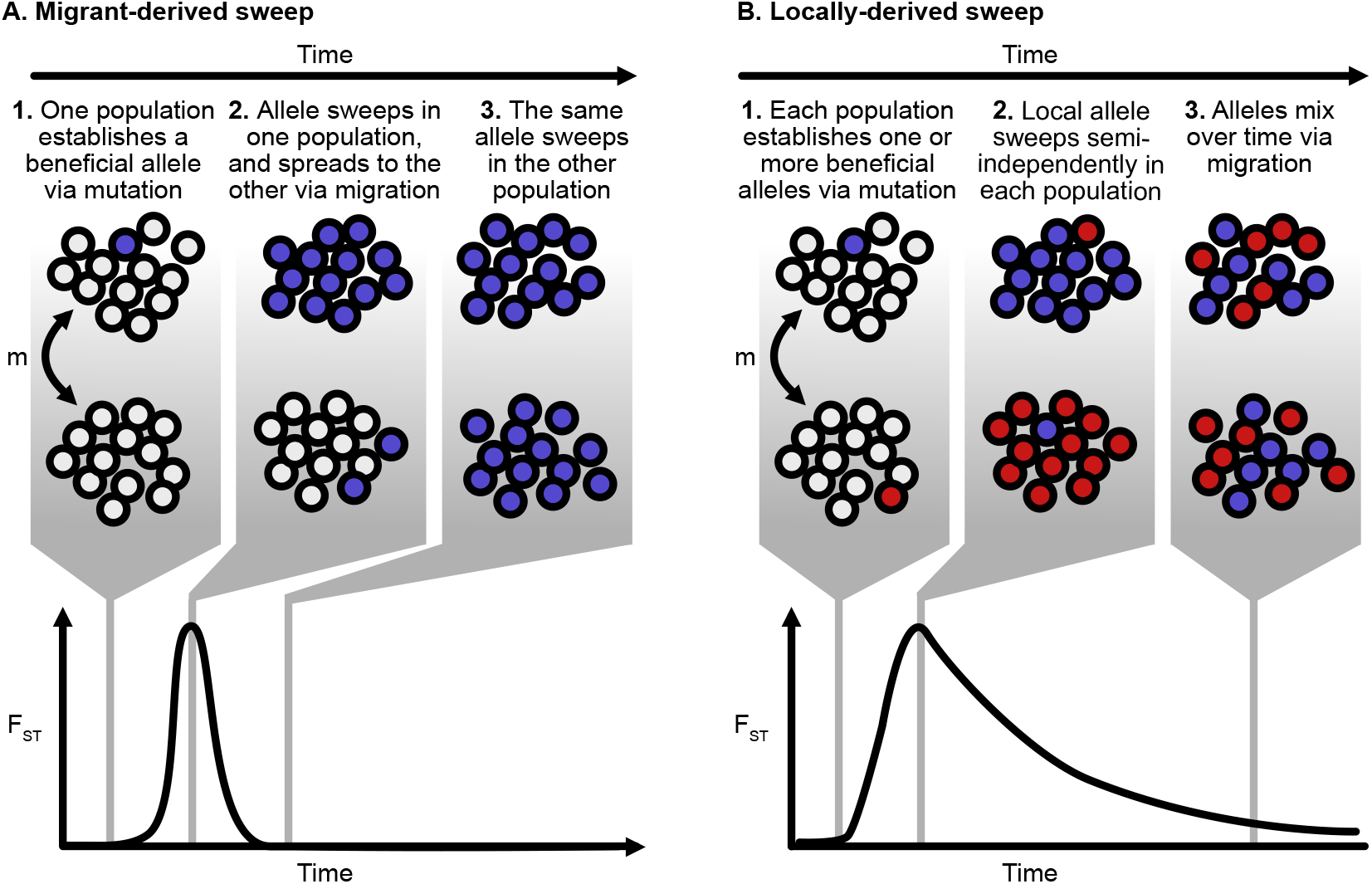
Schematic of allele frequencies and *F_ST_* across populations adapting in parallel. **A.** In a migrant-derived sweep, one population generates a beneficial allele via mutation significantly before the other population. The allele sweeps locally in its population of origin, and arrives in the alternative population due to migration before this population can fix its own variant. Here, the increase in *F_ST_* is due to divergent frequencies of the wildtype and derived allele, with the alternative population having a high frequency of the wildtype allele while the focal subpopulation has a high frequency of the derived allele. Migrant-derived sweeps have a short, symmetric spike in *F_ST_*. **B.** In a locally-derived sweep, each population fixes its own variant (shown in red and blue) which increases in frequency locally. Over time, migration equilibrates the allele frequencies among the two populations. The increase in *F_ST_* is due to divergent frequencies among different derived alleles across populations, and *F_ST_* has a long-tailed, asymmetric trajectory over time.

In a “migrant-derived sweep” (Figure 2A), a beneficial mutation sweeps in one subpopulation and then spreads to the second subpopulation via migration. The mutation then sweeps in the second subpopulation. Population differentiation increases due to different frequencies of the same allele (or alleles) across subpopulations, and then disappears when the sweep completes in the second subpopulation. This case, which has been described previously in [22] and [23], is more likely when the influx of beneficial mutations is limited (i.e., *θ* low).

When the population mutation rate is high, another path is also possible (Figure 2B). If many beneficial mutations enter the population each generation, then each subpopulation may produce its own beneficial variant (Figure 2B1.) which begins to rise in frequency locally (Figure 2B2). Population differentiation (as measured by *F_ST_*) increases due to sweeps of different alleles concurrently. However, over time, migration equilibrates the allele frequencies across the two populations (Figure 2B3), and *F_ST_* returns to longterm equilibrium levels. As we will explore later, the initial increase in differentiation and the rate of equilibration can be informative for the speed of migration.

The shape of the *F_ST_* trajectory over time differs substantially between the two cases. In locally-derived sweeps, selective sweeps drive the initial increase of *F_ST_* and migration erodes population differentiation over time. In the case of the migrant-derived sweeps, selective sweeps drive both the increase and decrease in *F_ST_*, because *F_ST_* decreases as the sweep completes in the second subpopulation (See Figure 2A). Migrant-derived sweeps are therefore substantially less informative than locally-derived sweeps about the rate of migration between two populations, because both the increase and decrease of *F_ST_* are on the timescale of selection.

Population genetic parameters (*s, m* and the population mutation rate *θ*) determine the relative probabilities of migrant- or locally-derived sweeps (see supplemental material). Because [23] describe migrant-derived sweep behavior in detail, we focus on locally-derived sweep behavior, although both types of sweeps occur in the model described below. Further, we believe a model of locally-derived beneficial mutations represents the more probable description of the dynamics in our motivating example of SHIV.

#### 2.1 Model

We consider a two-population island model with mutation, migration and selection. We model two haploid populations of size *N* with non-overlapping generations. Each population begins fixed with an identical wildtype allele (standing genetic variation is discussed below). Each generation, mutation introduces new beneficial mutations at a rate *Nμ* at a single locus, and selection increases the frequency of those mutations with strength 1 + *s* relative to wildtype (*s* > 0). All beneficial mutations are neutral with respect to each other, and no individual can carry multiple mutations. This describes loss of function mutations where subsequent mutations in the same pathway have no effect and follows the modeling assumptions of [24]. Each generation, a proportion of the population *m* migrates symmetrically between the two subpopulations. Therefore, *Nm* = *M* individuals migrate in each population per generation.

We investigate first the conditions under which the subpopulations will differentiate, as measured by the population differentiation statistic *F_ST_*.

Figure 3 shows a collection of stochastic simulations. Each colored region represents the spread of an allele in the two large populations experiencing strong positive selection (*N* = 10^5^, *s* = 1). Red alleles originate in population A and blue alleles originate in population B. When the migration rate is low, each population fixes independently its own set of alleles (see *M* = 10^−1^ through *M* = 100). When the migration rate is high, the populations are indistinguishable and represent a mix of alleles originating in both subpopulations (i.e., *M* = 10^4^). Among intermediate migration rates, however, more complicated dynamics arise. The population differentiation statistic *F_ST_* initially rapidly increases and then slowly decreases. The nonmonotonicity of these trajectories can be understood by considering three relevant timescales: 1) the relative forces of selection and migration during the sweep, 2) the relative forces of migration and drift following fixation, and 3) the relative timescales of migration and the sampling.

**Figure 3.**
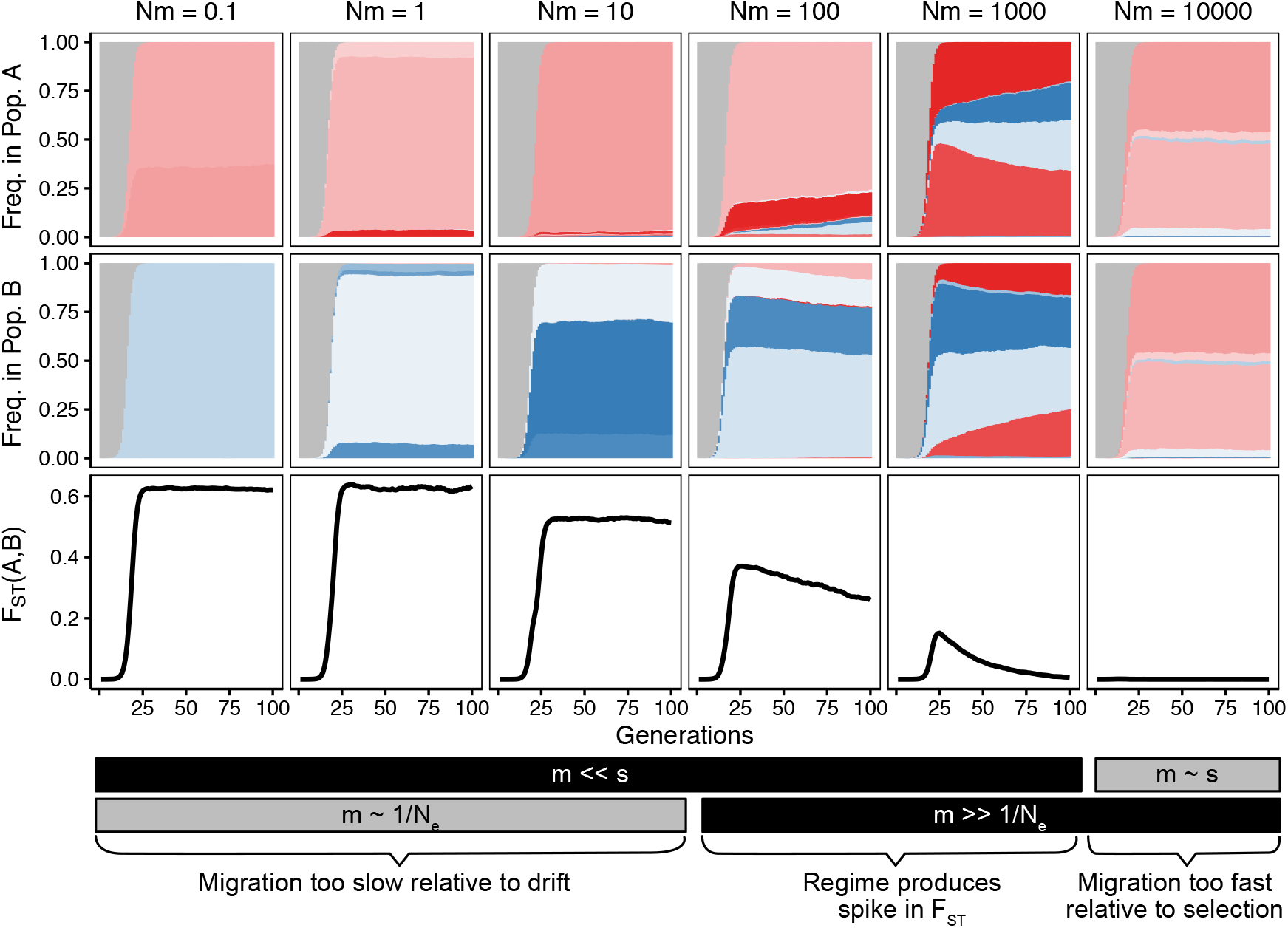
Patterns of allele frequencies and *F_ST_* across populations adapting in parallel. Beneficial alleles originating in populations A and B (colored red and blue respectively) increase in frequency over the wildtype allele (in grey) creating patterns of differentiation in *F_ST_* shown on the bottom row. The patterns of differentiation are dependent on the migration rate connecting the two populations (*M* = *Nm* shown in columns, *s* = 1, *N* = 10^5^, *m* varies as listed in column header.)

The initial dynamics are determined by comparing the forces of migration and selection during the sweep. If selection overwhelms migration (*s* ≫ *m*), each population at least partially fixes its own variant and *F_ST_* increases rapidly (here, *M* ≤ 10^3^). Alleles spread within subpopulations faster than migration equilibrates these alleles across subpopulations. However, if migration is too fast compared to selection, little differentiation occurs at any point in the sweep (*M* = 10^4^).

The subsequent decrease in *F_ST_* depends on the relative timescales of migration and both genetic drift and sampling post-sweep. In our model, all alleles share identical selective benefit and no secondary mutations occur on the background of the first mutation (although the first mutation can happen on the wildtype background multiple times). Therefore, after loss of the wildtype allele, all alleles are selectively neutral with respect to each other, and solely migration and drift govern the changes in their frequencies. *F_ST_* between the two populations will ultimately decrease to near the equilibrium predicted by Wright (*F_ST_* = 1/(1 + 4*Nm*)), but the rate of that equilibration depends on the migration. If migration is sufficiently fast compared to drift (i.e., *m* ≫ 1/*N_e_*), *F_ST_* will decrease faster than drift will move alleles within subpopulations. However, if migration is comparable to drift, the process will take significantly longer (on the order of *N_e_* generations). Whether this decrease is observed depends on the time frame of population sampling, although this paper will be restricted to relatively short sampling periods (i.e., 100 generations).

These observations suggest that in adapting populations, distinctive patterns emerge depending on the relative values of migration (*m*), selection (*s*), genetic drift (1/*N_e_*) and the sampling timescale. By examining these patterns over short timescales in genetic data, we may be able to estimate the values of *s, m* and *N* relative to each other in natural populations. This estimation will be a primary goal of the paper.

An analog of Figure 3 with a lower population mutation rate (*θ* = 0.5) is shown in Figure S1, and displays example patterns of *F_ST_* in parameter regimes where migrant-derived sweeps are prevalent. See supplemental text for discussion concerning these sweeps.

If a similar picture is produced using neutral alleles, only the lowest migration rates produce differentiable patterns and elevated *F_ST_* (Figure S2), and it takes much longer for this differentiation to accrue (i.e., on the order of *N_e_* generations).

The dynamic patterns of *F_ST_* under strong selection and migration suggest sweeping alleles across populations can provide insight into migration that is too fast for neutral alleles. As in the neutral case, when migration is too slow compared to the rate of change in the population we can only say that migration rate belongs to a particular, slow regime. We encounter a similar situation when migration is too fast compared to selection. However, in the intermediate cases when selection and migration are on sufficiently similar timescales, more precise estimation appears possible. By observing alleles moving at different speeds due to different strengths of selection, we can move the boundaries of the bins and estimate more precisely migration at different speeds.

This example also illustrates several additional practical points. 1) Because dynamical patterns in *F_ST_* are apparent when sweeping alleles are captured at multiple points in their trajectories, this approach depends on the availability of time series data. 2) Because differences in the allele frequencies must be measurably different, sampling depth will affect whether or not these signatures are present.

Before dynamical patterns of population differentiation can be used to perform parameter estimation in a stochastic model, we first examine patterns of *F_ST_* under strong migration and selection in a deterministic setting. Specifically, we will explore analytically the parameter regimes that create transient and long-term population differentiation driven by locally-derived sweeps. We will see later that the population parameter regimes that create nonmonotonic signatures of population differentiation correspond to those in which we can successfully perform parameter estimation using approximate Bayesian computation.

#### 2.2 Analytical approximation

To study locally-derived sweeps, we consider the case in which each of two populations has a single copy of a local allele at the same locus (i.e., frequency *f*(0) = 1/*N*) simultaneously. We ignore all subsequent mutation. Because the populations are symmetric, without loss of generality, we can consider the frequencies of a single subpopulation and determine what are the relative frequencies of the local and non-local allele at a time *t*(*f_l_*(*t*) and *f_nl_*(*t*), respectively). We consider the total number of derived alleles *f*(*t*) = *f_nl_*(*t*) + *f_l_*(*t*) and assume that *f*(*t*) grows logistically.

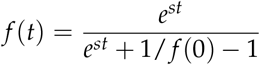

A differential equation describes the change in frequency of the non-local allele, *f_nl_*(*t*):

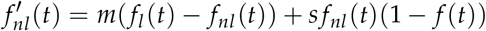

We define *F*(*t*) = *f_nl_*(*t*)/*f*(*t*), the frequency of the nonlocal allele among all derived alleles. We solve to find

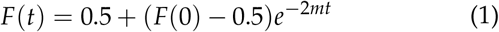

Closed form descriptions of *f_nl_*(*t*) and *f_l_*(*t*) are given as follows:

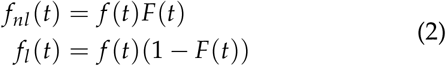

If we assume that each population starts with its own allele at arbitrarily low frequency (i.e., *f_l_*(0) = 1/*N* and *f_nl_*(0) = 0), we can describe the dynamics of the two alleles over time via *F_ST_*:

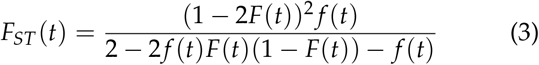

The predicted *F_ST_* trajectories match well the trajectories generated by simulations showing locally-derived sweeps (Figure 4). Because multiple circulating alleles constrain the possible values of *F_ST_* [25], all alleles originating from the same population from simulation are collapsed into a single allele so that the *F_ST_* magnitudes are also comparable.

**Figure 4.**
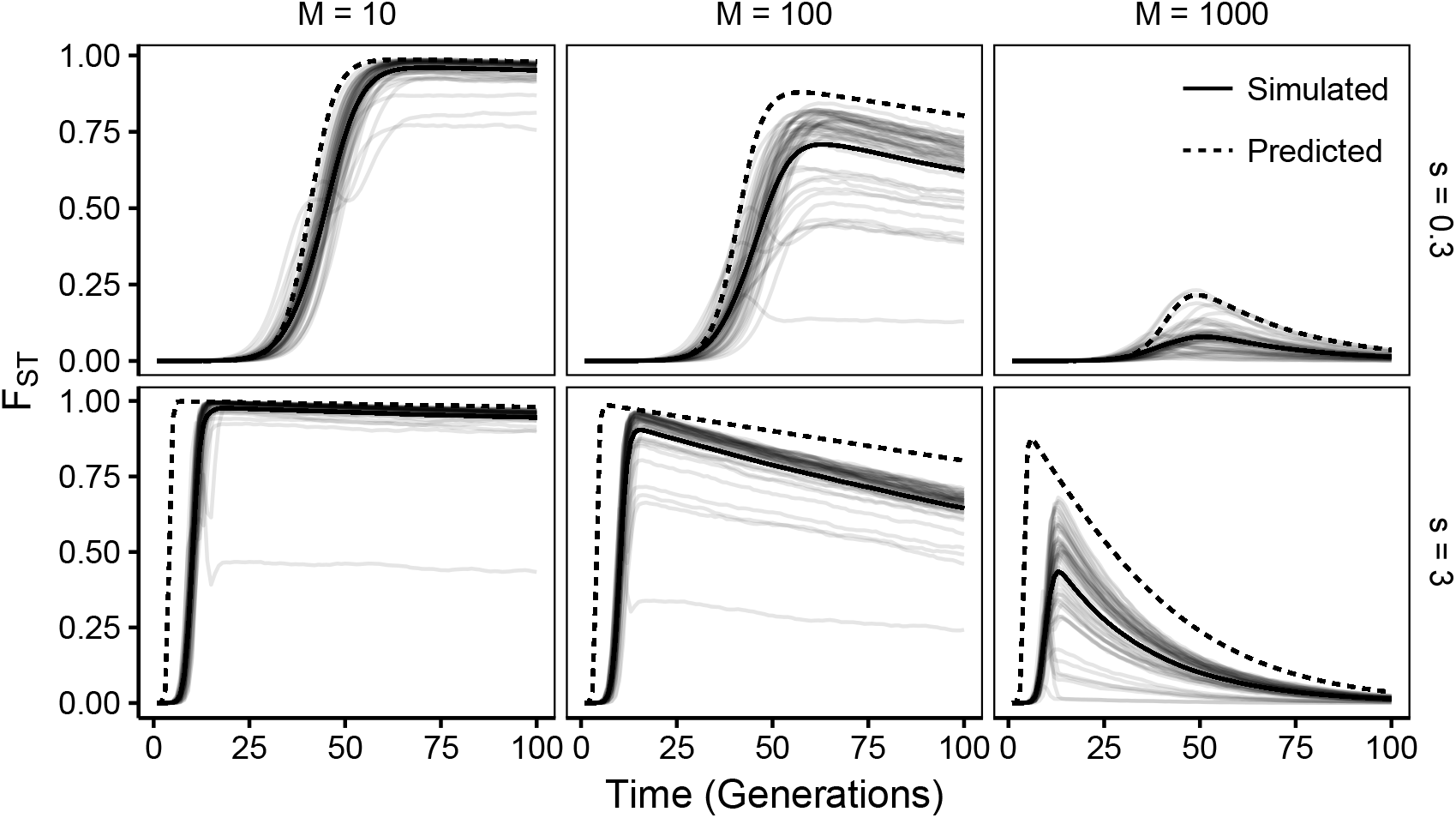
Simulated patterns of *F_ST_* over time share features with analytical predictions. Forward stochastic simulations of *F_ST_* between two populations undergoing parallel selection of strength *s* and connected by *M* variants per generation for a given set of parameters are shown in light, solid lines (100 replicates). All alleles originating in a single subpopulation are collapsed together for the purpose of computing *F_ST_*. The median trajectory is shown in a dark solid line. Analytical predictions (equation 3) are shown in a dashed line. (*N* = 10^5^).

This equation also allows us to quantify the generation at which *F_ST_* will be maximized between the two subpopulations (*t_max_*), and its maximum value (*F_ST_*(*t_max_*)):

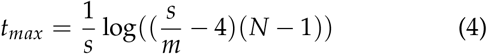

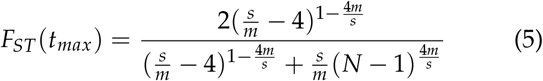

From these equations, we can determine that the waiting time until the population reaches maximal *F_ST_* is dominated by a term inversely proportional to *s*. However, the maximal *F_ST_* at this point is determined by the ratio of *s/m*. The strength of selection controls largely when the *F_ST_* spike will happen, but the ratio of *s/m* governs the magnitude of the signal, with stronger selection relative to migration creating more pronounced spikes as a non-linear function of that proportion. Note that this equation is undefined when selection is not sufficiently stronger than migration (*s* < 4*m*).

Below we will also use these equations to quantify the parameter regimes in which *F_ST_* will show non-monotonic patterns by evaluating when *F_ST_*(*t_max_*) will be elevated sufficiently for it to be observed at a given depth, and also when *F_ST_* will decline appreciably over a relevant timescale by evaluating equation 3 some number of generations after *t_max_*.

### 3. Using patterns of population differentiation for parameter estimation using approximate Bayesian computation

To translate these expected patterns into parameter estimates, we employ approximate Bayesian computation (ABC) [26, 27] because exact likelihood formulations cannot be worked out under complex demographic scenarios. This provides a framework for interpreting allele frequency trajectories in the context of underlying parameter regimes. Briefly, ABC works through simulating data under parameters drawn from a prior and then comparing the simulated data to observed data. Parameters that generate simulated data similar to observed data form posteriors. We do not intend this as a claim for the best way to estimate migration using selected alleles, but as a demonstration of extractable signal.

We begin by demonstrating that migration information can be estimated from observed allele frequencies generated from our stochastic simulations using allele frequency differences in these connected populations over time. Recently, many methods have built up around the idea of estimating selection strength from allele frequency data in a single population [28, 29? ?]. By leveraging this information across multiple populations, we can learn about selection and migration simultaneously. In particular, we imagine that we have allele frequency samples of *n* genomes before and after a sweep, and at some time point *T* generations later (here: *n* = 100, *T* = 30). These time points are chosen to match the Simian-HIV motivating example, but we provide a much more thorough analysis of how sampling time affects this approach below.

Because we are tracking the frequencies of many alleles across multiple time points, we use summary statistics to characterize the observed dynamics. As noted in [30], the accuracy of ABC can be boosted when summary statistics are optimized separately for estimating different parameters, especially when there are only weak interactions between pairs of parameters. To jointly estimate *s, N* and *m*, we selected summary statistics that were useful for fitting the migration rate separately from *s* and *N*. We first performed a single round of ABC to estimate posteriors over *s* and *N* with these summaries. Then, we performed a second round of ABC to estimate *m* while restricting the priors over *s* and *N* to the posteriors from the first round fit (see Materials and Methods). We also fit *s* and *N* separately from each other and did not find a significant improvement in method performance (data not shown). To summarize information about *N*, we found the most likely *θ* using Ewens’ Sampling Formula for the combined allele frequencies at each sampled time point pooled across the populations. To summarize information about *s*, we assumed beneficial allele frequency 10^−5^ for a beneficial mutation at time 0 (which is a conservatively low frequency for intra-host standing genetic variation [31, 32]) and determined the parameter *s* producing the logistic growth curve that minimized mean squared error from the observed data. To compute *M*, we used four parameters at each time point: *F_ST_*, 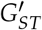, the difference in heterozygosities between the two subpopulations and the number of shared alleles at any frequency. Reasoning and further details concerning these summary statistics can be found in the Materials and Methods.

**Figure 5.**
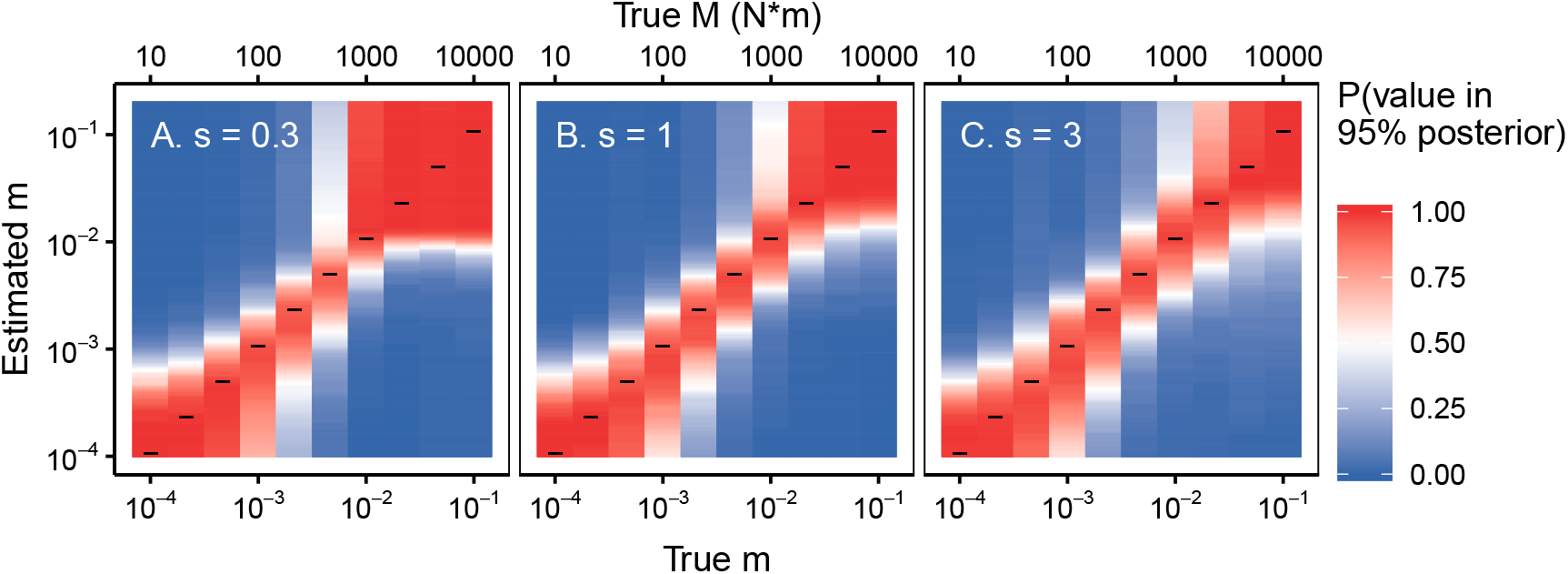
ABC procedure re-estimates true *M* values in simulated data. 200 simulations are run for each true value of *m* on the *x*-axis and 200 posteriors are produced. We plot the proportion of 95% posteriors for a given true *m* (shown on the *x*-axis) that include the various values of estimated *m* (shown on the *y*-axis). Values that are more red indicate that the posteriors successfully capture the test ‘Estimated *m*’ value on the *y*-axis whereas blue values indicate that few posteriors include the *y*-axis value. Black horizontal lines indicate where the true value matches the *y*-axis ‘Estimated *m*.’ Simulations are generated across a gradient of values for *m* and *s* (*m* ∈ (10^−4^ – 10^−1^), *s* = (0.3,1,3), sampling at generations (5, *T* = fixation, *T* + 30,100), *N* = 10^5^).

#### 3.1 Migration estimation depends on *s*

To validate our ABC approach, we first simulated data under known parameters and then used our approach to reestimate migration rates. We ran 200 forward simulations for each set of parameters and examined the resulting 95% posteriors over *m* to quantify the proportion of posteriors that contain the true *m* (Figure S4). See Materials and Methods for full simulation details. We also quantify the probability that the posteriors contain incorrect values of *m* at varying distances away from the truth. We find the 95% posteriors contain the true migration rate with high probability (Figure S4, indicated by the clustering of red around the *x* = *y* line) and do not contain migration rates far away from the truth (indicated by off *x* = *y*).

There are instructive exceptions. First, when the migration rate is low, we have little power to distinguish values of *m* in this range. Low *m* results in near total differentiation between the two subpopulations (see Figure 3, *M* < 1000), so it is therefore unsurprising that we cannot predict the migration rate beyond bounding it. Similarly, this method cannot differentiate among very high migration rates. In this instance, the populations are entirely panmictic, and there is little signal to uncover (See Figure 3, *M* = 10^4^). Although within these regimes where allele frequencies from the two subpopulations appear panmictic or independent we have no ability to estimate migration rates specifically, this does allow us to bound fast migration rates much more precisely than is possible with neutral alleles. For example, if we were to observe identical equilibrium neutral allele frequencies among two subpopulations, we might conclude that *M* > 1. However, if we observed identical allele frequencies over time subject to strong selection, this would suggest that *M* ≥ 10^4^.

The boundaries at which we enter these regimes of independence and panmixia depend on the strength of selection. We see that *s* = 0.3, *M* = 1000 results in apparent panmixia in many instances. However, when *s* = 3, our method still accurately estimates migration rates when *M* = 1000 (Figure S4). We test this more directly by computing the log MSE between the posteriors and the true *m* for a wide variety of selection strengths and migration rates (Figure 6A, see Materials and Methods for a detailed description of the log MSE). Consistent with the analysis in Figure S4, when migration is too high or too low relative to selection, the posterior is far from the truth. (Note, the relatively low MSE between the truth and the posterior among high values of *m* is driven by the limits on the prior, which is bounded above by *m* = 0.5). However, the upper boundary at which the performance deteriorates increases with selection strength. Put another way, as selection strength increases, so do the rates of *m* that can be accurately estimated.

**Figure 6.**
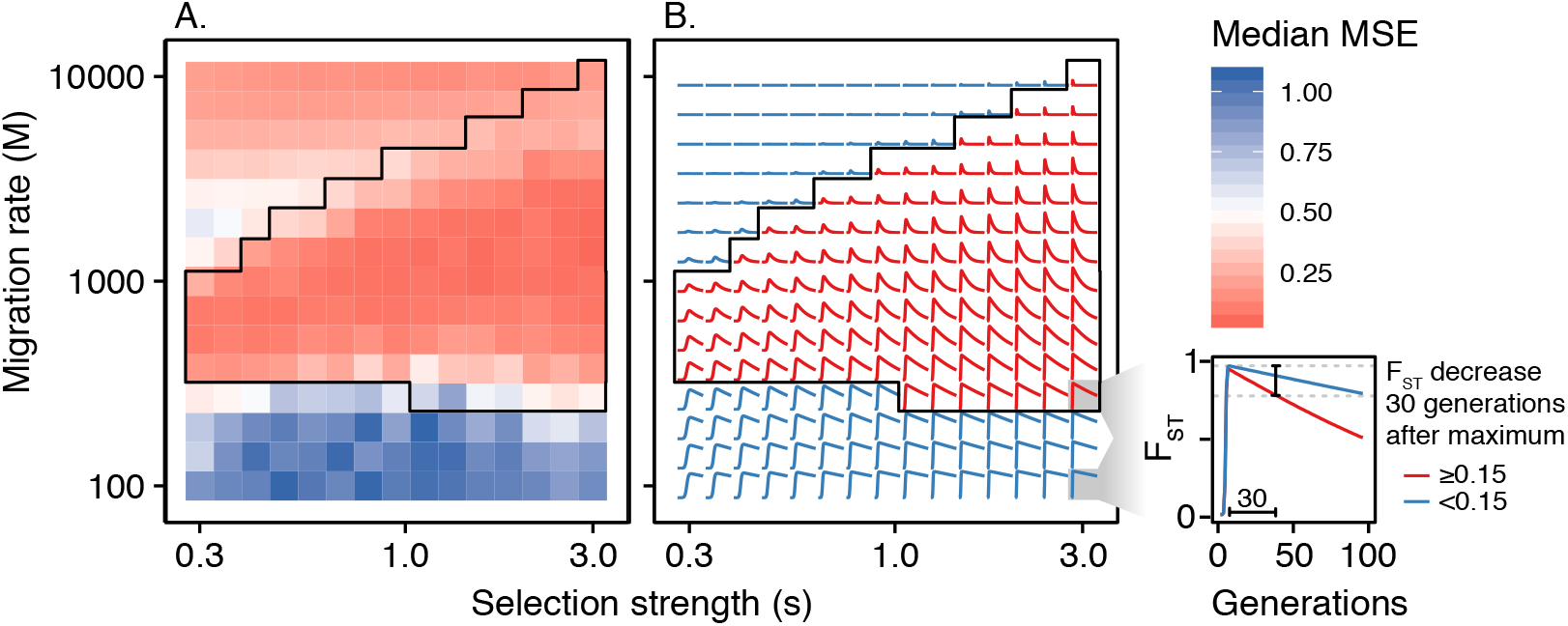
Better estimation of higher migration rates among simulations with higher selection strength. For a fixed *N*, higher *s* values result in lower log MSEs (indicating successful estimation) for higher values of *M*. Each tile represents the median log-MSE for 100 trials computed with a given *m* and *s*. (*N* = 10^5^, sampling at generations 5, *T* = fixation, *T* + 30). **B.** Predictions of the dynamics of *F_ST_* over 100 generations under a variety of parameters for *s* and *m*. Each square in **A.** corresponds to the modeled dynamics in **B.** The parameter combinations predicted to lead to a 0.15 unit decrease in *F_ST_* after 30 generations (red) correspond to the parameter regimes where migration can be estimated with low MSE. The area circumscribed in black highlights this region in both **A** and **B**.

These results are predicted by our analytical model of locally-derived sweeps. The parameter combinations that show a noticeable non-monotonic pattern in *F_ST_* over 100 generations (i.e., *F_ST_* reaches a value greater than or equal to 0.15 followed by a decrease in *F_ST_* > 0.15 within 30 generations of *t_max_*) are similar to those that can be estimated using the ABC approach (6B). This suggests that a primary signal identified by our ABC approach is a non-monotonic pattern of *F_ST_* over time. However, because the performance of the ABC approach deteriorates as other summary statistics are removed (data not shown), this is likely just one of several important signals for estimation.

#### 3.2 Practical considerations in estimating *m* from data

The interaction between migration and selection is not the only factor that impacts our estimation. We find that the performance improves when the sampling is deeper, reflecting more accurate estimation of allele frequency differences (Figure S3). We also find that the choice of sampling time points matters for the estimation of specific migration rates. We assume that samples are taken before and after a sweep, and then at an additional time point *T* generations later. When *T* is small, performance is worse, presumably because migration has had less time to influence allele frequencies (Figure S4). When adding a fourth time point at generation 100, the method is fairly insensitive to the placement of the third time point (Figure S5).

We also find that specific parameters of the ABC are important, namely the tolerance of accepted matches. We find that as tolerance becomes more stringent, the posteriors become narrower (Figure S6A). However, as the resulting posterior decreases in size due to lower tolerance, there is a decreased probability of capturing the true value in the posterior (Figure S6B). Nevertheless, decreasing the tolerance improves the matching of posteriors and the truth (Figure S6C), and we therefore choose a low tolerance (*tol* = 0.001) for our analyses.

#### 3.3 Application to real data

In the previous section, we have established that we can estimate population genetic parameters using a multi-step ABC method. In this section, we apply this method to real data that matches the structure of our ABC method [17] in order to estimate viral migration rates between pairs of different organs (blood plasma, lymph nodes, gut) of a Simian-HIV infected pigtailed macaque sampled over time (Figure 1). The importance of estimating such rates has been previously investigated (although in the absence of selection) [33]. The allele frequencies shown in Figure 1 are simplified trajectories of the full data to make it more comparable to our model. For full data processing choices (and fits when those choices are modified), see the Materials and Methods section.

We find that estimates of *m* between the plasma and lymph node are very high: The distribution is flat above 1% of the population migrating each generation (Figure 7, Table 1). The flatness of this distribution suggests that these subpopulations exist in the regime in which migration moves alleles faster than selection, and we find little differentiation that cannot be explained by sampling error (consistent with findings of panmixia in [17]). Although a more specific migration rate cannot be estimated, posteriors suggest a migration rate above *m* ≥ 0.001.

**Figure 7.**
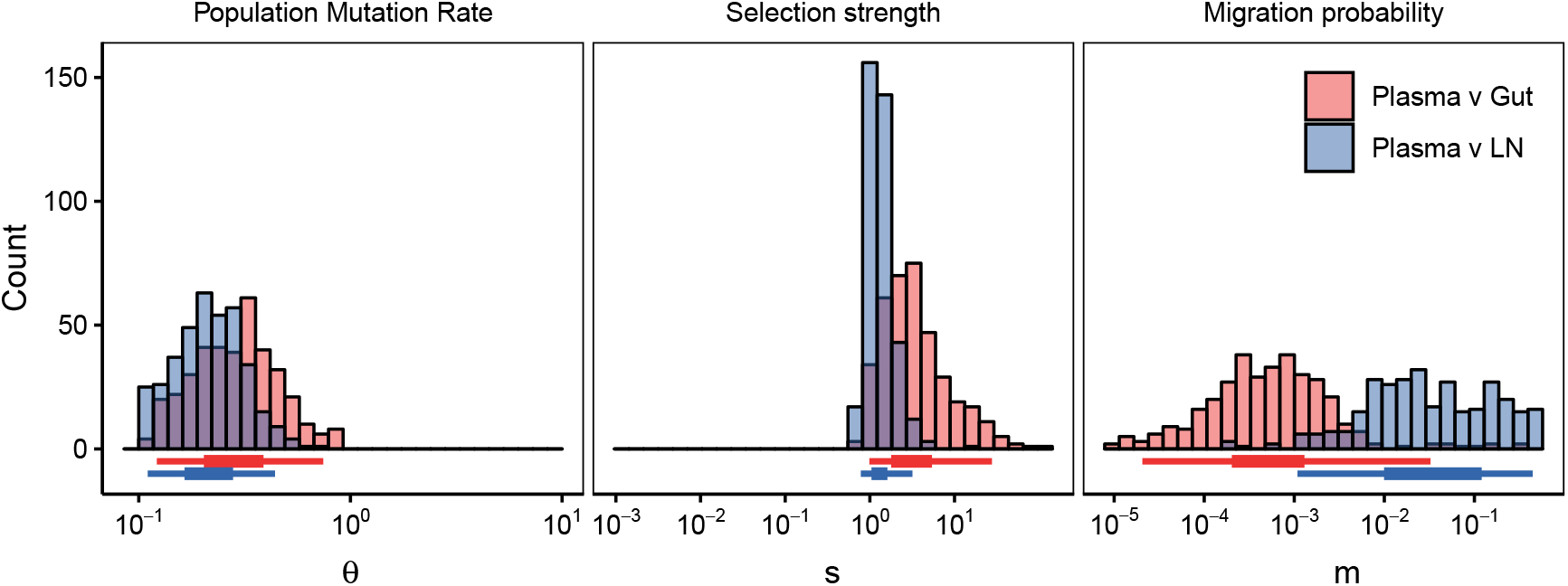
Estimation of population parameter rates from intra-patient Simian-HIV data sampled from different subpopulations. The top row shows diagrams of drug resistant haplotypes spreading in different subpopulations over time sampled at generations 7, 21,49 and 98 in the gut, lymph node and blood plasma. Each color represents a distinct lineage separated by at least one mutation. The resulting posteriors are given for the ABC procedure for comparisons between the plasma and gut (red) and plasma and lymph node (blue). 95% posteriors for these distributions are shown in Table 1.

**Table 1.** Medians and 95% posteriors for parameter estimation in intra-macaque Simian-HIV populations.

| | $m$ | $M$ | $\theta$ | $s$ |
| --- | --- | --- | --- | --- |
| Plasma v Gut | 0.000528<br>(2.06e-05,0.0326) | 15.5<br>(0.602,953) | 3<br>(0.987,27.7) | 0.293<br>(0.122,0.744) |
| Plasma v Lymph Node | 0.0271<br>(0.00109,0.444) | 573<br>(23,9400) | 1.24<br>(0.783,3.19) | 0.212<br>(0.11,0.44) |

However, the migration rate between the plasma and the gut is unimodal centered around 0.05% of the population migrating each generation (Figure 7, Table 1) - half of the low end of the posterior between the lymph node and the plasma, yet higher than what would produce elevated *F_ST_* in drifting alleles at equilibrium (although *N_e_m* = 1 falls within the 95% posterior). These differences between the connectivity between the plasma and gut or lymph node are also ordinally reasonable, as we might expect the circulatory system to be more connected to the lymphatic system than a mucosal tissue.

We estimate similar selection strength and population mutation rates across subpopulation comparisons (see Table 1). Across both comparisons, the selection strength is extremely strong (*s* ≈ 1.24 - 3), which is necessary to explain the almost complete sweeps observed in the first 20 generations. The population mutation rate is sufficiently high that soft sweeps are likely [34]. That we see no significant differences between the population sizes makes intuitive sense given the consistency of sweep timing and diversity across the three populations. However, it is also possible that the dynamics are driven by the larger subpopulation, resulting in comparisons that are independent of the secondary subpopulation.

Note, we make filtering choices to determine what is counted as a distinct haplotype (see Materials and Methods for full details). In the main text, we present intermediate filtering in which alleles observed in at least two copies before drug resistance sweeps (>70%) are included. We can repeat the analysis for alleles observed in at least one copy (Figure S7) or at least five copies (Figure S8). Different haplotype thresholds unsurprisingly alter estimates of *θ*, but posteriors are robust for estimates of *s* and *m*, likely due to the factorization procedure (Figures S9 and S10). Posteriors for all three allele frequency cutoffs are listed in Table S1.

## Discussion

*F_ST_* is a widely used metric across ecology and evolutionary biology. Although *F_ST_* is problematic in a variety of ways, its ubiquity means that it is worth understanding in detail. In particular, many studies use Wright’s characterization of the relationship between the migration and population differentiation as a prescription for estimating migration rates. Problems with this approach have been previously discussed [16], but here we show that on short timescales, *F_ST_* can depart drastically from the neutral equilibrium due to strong perturbations. We focus on how strong selection can distort *F_ST_*, but this holds more generally true for populations perturbed by a variety of ecological scenarios (for example, population bottlenecks).

That *F_ST_* behaves dynamically over very short timescales has important implications for interpreting population differentiation data. In particular, data sampled at a particular time point can be very misleading in determining population dynamics. Even with constant migration rates, samples at different time points could lead one to conclude that migration is either prevalent or rare, and migration rates much larger than *M* = 1 can lead to substantial population differentiation transiently. This suggests that merely observing elevated *F_ST_* is not a sufficient condition for diagnosing a low-migration system and understanding non-equilibrium dynamics of these statistics is important for analyzing data. Proper accounting for the covariance among selected alleles can help disentangle the signatures left by combined selection and migration [21].

Although migrant-derived sweeps have been described previously, locally-derived sweeps can help explain the apparent “softening” of hard sweeps in HIV, which initially show only a single beneficial haplotype, yet later multiple haplotypes appear [35, 24, 34, 36]. If each subpopulation fixes its own beneficial variant, which later mix, a locally hard sweep can appear to become soft. This is in contrast to the previously described “hardening” of soft sweeps, which can occur when demography causes only a single beneficial haplotype of a soft sweep to fix [37].

To understand the conditions in which *F_ST_* is elevated over short timescales, we explore an analytical model of population differentiation under strong migration, selection and mutation. In time series data, we find a variety of dynamics of elevated *F_ST_*: if selection is stronger than migration (*s* ≫ *m*), *F_ST_* will increase over short timescales. If migration is faster than drift (*m* ≫ 1/*N_e_*) and the sampling timescale, *F_ST_* will quickly decrease after the sweep. When both conditions are met, characteristic non-monotonic patterns in *F_ST_* emerge over experimentally tractable timescales. We introduce a modeling framework to further explore this diversity of patterns.

That many patterns emerge with different parameter combinations of *Nμ, s* and *m* suggests that measurements of population differentiation over time could provide information about migration rates much larger than those normally considered and therefore an opportunity to measure such rates. Because the patterns depend on the relative strengths of these population genetic forces, we expect that these patterns need to be estimated jointly. This is problematic because we know that many population genetic forces act on composite parameters (for example, *N_s_*, and *θ*). However, we also observe that different types of statistics can give us information about different parameters, as some parameters have a strong effect on one statistic and a smaller effect on another. For example, changes in *s* cause drastic changes in the rate of change of allele frequencies in a population, but provide little difference in the number of circulating alleles (i.e., the softness of the sweep). This provides an opportunity to “factorize” the different parameter estimation into relevant statistics (similar to what is described in [30]).

Having learned that we can disentangle information about different parameters using different types of statistics, we develop an iterated-ABC framework that estimates *N_e_, s* and *m* jointly. We explore the regimes where estimation is accurate, and find that its accuracy can be predicted from a simple analytical model. In the parameter regimes where estimation is not accurate, we can bound migration rates above or below certain values by comparing the rate of migration to the processes of drift and selection. These broad regimes of estimation suggest that we can estimate a wide range of migration rates that we might expect to be important among pathogen populations.

As an example of the approach, we estimate migration rates between viral populations inhabiting different organs in a Simian-HIV infected macaque treated with drugs. Consistent with the findings of the original study which used several metrics of population differentiation but did not attempt to estimate rates, we find differential connectivity between compartments of the body. In particular, we find that mucosal tissues (i.e., the gut) have lower migration rates into the blood than the lymph nodes do. However, using our ABC framework, we can make this description of the intra-patient environment quantitative as opposed to simply qualitative. We estimate per-virus migration rates that can be used in future modeling studies.

Our results also provide guidance for experimental design, particularly in non-mutation limited systems with strong selection and potentially high migration rates such as pathogen evolution, and pesticide and herbicide resistance. In particular, we find that the most useful way to glean information about migration rates from data is to include a temporal component to sampling. For migration estimation, it is considerably more useful to have three time points of 30 sequences than a single time point of 90 sequences. It is particularly useful to sample during or shortly after a selective sweep. From a practical perspective, the timing of such sweeps can be determined by phenotypic shifts in the population. If only a single time point directly after the sweep is available, this can provide information about whether migration is slower than selection. However, if an additional later time point is available, this potentially also gives information about whether migration is faster than drift. Having samples at both of these times provides both upper and lower bounds for high migration rates.

Although our method provides an avenue to learn about high migration rates, these estimates should be taken with same credulity that we apply to *F_ST_* under Wright’s neutral island population model [38]. In natural systems, many of these model assumptions are likely broken and can also lead to elevated population differentiation in adapting populations. In the context of intra-patient virus evolution, many of these assumptions are likely to be violated. Drugs may reach different tissues with variable concentrations, creating weaker or stronger selective pressure. The virus could be adapting locally to different tissues, creating mutations that are deleterious in other parts of the body. We also know that viral population sizes can differ greatly between tissues.

Despite the violation of model assumptions in natural systems, this paper provides a null model for observable patterns with high migration and selection. We show that transiently high differentiation is possible in adapting populations without needing to invoke common explanations such as local adaptation or low migration rates. In the future, if evidence of local adaptation between populations is proposed, investigators will need to further show that such patterns cannot be caused with the same selective pressure across subpopulations. Complex phenomena may not necessarily need to be invoked to explain the data, and it remains to be seen how we can detect these phenomena given the complexity that we see even under the simplest of null models with uniform selection. For example, in the Simian-HIV data, different selection strengths in different compartments (for example, from differences in penetration depth) are not necessary to explain the observed patterns. We observe rapid adaptation in all three compartments with different mutations picked up. Alternative explanations are certainly possible with more complex scenarios, in particular with local adaptation, but in certain instances, we may not need complicated mechanisms.

An additional problem is that in non-mutation limited systems, not only do multiple mutations arise simultaneously, but populations quickly acquire double and triple mutants. It is therefore important that in future studies, we consider how these patterns of population differentiation might appear on a traveling wave of beneficial fitness effects, similar to what has been done in single populations [39] and spatially structured populations with lower migration rates [24].

To understand very rapid processes (such as fast migration) or the forces governing the population in its current state, it is insufficient to look at dynamics averaged out over long time periods. Selection provides an avenue to observe the current population state in time that neutral processes can miss. Using long term metrics (like comparing the rate of migration to the rate of drift) to investigate how much migration is happening in the moment invites numerous misinterpretations. When studying migration, as when studying all population processes, the correct timescale must be carefully considered.

## Materials and Methods

### Simulation

Two populations of equal size *N* are instantiated with no standing genetic variation, under the assumption that beneficial mutations are costly before the change in environment. For Simian-HIV, this corresponds to fitness cost of DRMs in the absence of a given drug. Each generations, each population draws a Poisson distributed number of mutations (*λ* = *N* * 10^−5^) which land on wildtype backgrounds. All beneficial mutations confer identical fitness benefit 1 + *s* relative to wildtype. Following [24], we allow only a single beneficial mutation per individual, so once a sweep is complete (no wildtype individuals remaining), no additional mutations enter the population.

### Approximate Bayesian Computation

We performed a two-step approximate Bayesian computation procedure in which we simulated 3 × 10^6^ forward trajectories with uniform *1og*_10_ priors where *m* ∈ (10^−5^,5 × 10^−1^), *s* ∈ (10^−1^,10^2^), and *θ* ∈ (10^−1^,10^1^). We constricted our *θ* and *s* posteriors to values that could produce a sweep in the first 100 generations.

We used rejection sampling, which accepts a certain percentage of trials (given by the tolerance) that minimize the Euclidean distance between the observed summary statistics and those summary statistics generated by the prior as implemented in the R package ‘abc’ [40]. The parameter combinations that result in the lowest distances form the posterior. We first perform ABC to estimate the posteriors for *θ* and *s*, using targeted summary statistics as discussed in [30].

For *θ* we use the best fit *θ* under Ewens’ Sampling Formula for when the populations are combined at each time point. Note, combining two population allele frequencies with limited migration overestimates the value of *θ* (i.e., increases diversity relative to a single population), but this does not affect our procedure because the best fit *θ* is used only as a summary statistic.

To estimate *s*, we fit the observed frequency of the derived allele at the sampled time points to a logistic curve with y-intercept of 10^−5^ that minimized mean squared error from the observed data. In the cases of ties, we used the shallowest slope that could explain the data equally well.

From our first fit ABC, we get posteriors over *θ* and *s*. We then take the range of values within the 95% posterior for these two parameters and use those as a flat prior for a second fit (see Figure S11). For our second fit ABC, we use four summary statistics at each time point: *F_ST_*, 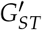, the absolute value of the difference in heterozygosities between the two subpopulations, and the number of variants shared at any frequency. For a schematic illustration of the two-step process, see Figure S11.

### Model evaluation

We determine whether our posteriors are providing useful estimates by comparing the posteriors to the true model parameter. The most useful posteriors will have small variance and be centered around the true values. However, using either one of these metrics will lead to misidentification. For example, if we evaluated a posterior using the proportion of the time the 95% posterior interval contains the true value, a completely uniform estimate across the posterior should contain the true value 100% of the time while offering no information about the estimated parameter. Alternatively, if we evaluated a posterior using the size of the 95% interval, a very narrow posterior far away from the true value would score highly but provide misleading information. To balance confidence and accuracy, we use the mean-squared error computed in log space (as our posteriors are uniform on a log-scale) to find the average difference between each of *n* points (*σ_i_*) in our estimated posterior and our true test parameter *x*.

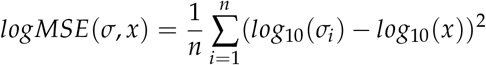

### Simian-HIV data

We analyze macaque T98133 from [17]. Briefly, the macaque was infected with RT-SHIVmne027, a simian immunodeficiency virus with an HIV-1 reverse transcriptase region and treated with FTC, a reverse transcriptase inhibitor. Drug resistance mutations to FTC occur within the reverse transcriptase region. The viral population was given 12 weeks to establish within the host before treatment began. Viral samples were collected from the gut, lymph node and blood plasma at 1, 3, 8 and 14 weeks after the onset of treatment. The authors performed single genome sequencing of the reverse transcriptase region, resulting in 800 bp regions containing relevant drug resistance mutations and complete linkage information and have approximately 30 sequences per time point and location.

Generations in HIV/SHIV are approximately 24 hours, so we translated weeks 1, 3, 8 and 14 to generations 7, 21, 56 and 98, which we rounded to generations 5, 20, 50 and 100.

We called derived haplotypes as unique if they contained a drug resistance mutation (M184V/I) and any other mutations at the initial time point at which drug resistance was observed (approximately generation 20) in at least *c* = 2 copies. Note, we replot Figures 1 and 7 with choices of *c* = 1 and *c* = 5 in Figures S7, S8, S9 and S10. 95% posteriors across the choices of *c* are also listed in Table S1. We excluded haplotypes appearing for the first time after generation 20, to avoid mutations occurring on the backgrounds of existing lineages, which our model does not consider. These later time point haplotypes were clustered back to the most common early time point haplotype with the minimal mutational distance. For example, if generation 20 contained three haplotypes with the mutations A, B and C: A, A+B and A+C, and the haplotype A+B+D was observed at generation 50, it was counted as haplotype A+B.

No additional known drug resistance mutations appeared after generation 20, although positive selection may still influence allele frequency trajectories.

When we compute *F_ST_* pairwise between anatomical compartments (as in Figure 1D), we treat each haplotype as equally different from every other haplotype, despite some sharing more alleles than others. For example, M184I and M184V+N255N+D177N and WT are all equally different from each other. We do this in line with our single mutation simulation model, in which all haplotypes are equally different from each other. Computing differentiation statistics that take into account conservation does not qualitatively change the patterns, and this question is considered in much more detail in [17]. This convention is also used in the computation of all other summary statistics.

### Code availability

All code to reproduce the above analyses can be found at https://github.com/affeder/fst_dynamics.

## Acknowledgements

We would like to acknowledge Ilana Arbisser, Jamie Blundell, Graham Coop, Jonathan Kang, Grant Kinsler, Chuan Li, Michael MacLaren, Rohan Mehta, Noah Rosenberg, and Lawrence Urichio for useful discussions on the content of the manuscript. This work was supported in part by the Miller Institute for Basic Research in Science at the University of California Berkeley (AFF), a Stanford Computational, Evolutionary and Human Genomics Trainee Fellowship (AFF) and ABI-1458059 (DAP).

## Supplemental Text

### Probability of locally-derived or migrant-derived sweeps

Based on the relative rates of migration, selection and mutation, we can predict whether we are likely to see sweeps driven primarily via migration or de novo mutation within subpopulations.

We assume subpopulation *A* has a derived allele at establishment frequency. We ask, for the second subpopulation *B*, what is the probability that the subpopulation acquires its first beneficial variant due to de novo mutation rather than migration from the other subpopulation. Note, this does not tell us which variant will ultimately sweep, as these variants can be lost before reaching establishment frequency, or can sweep together. However, we can build intuition on how migration, selection and mutation affect the probabilities of local- or migrant-derived behaviors.

The rate of alleles entering population *B* via mutation each generation is constant:

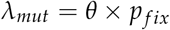

The rate of alleles entering population *B* via migration per generation from subpopulation *A* is

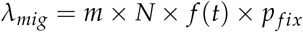

Where *f*(*t*) is the frequency of the derived allele in the subpopulation *A*. If we assume that *f*(*t*) is exponentially growing, then

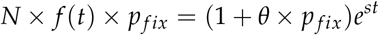

where *θ* × *p_fix_* accounts for subsequent mutation in *A*. Then, the probability that no derived alleles enter population *B* through time *t* is the probability of no mutations entering via mutation and no mutations entering via migration.

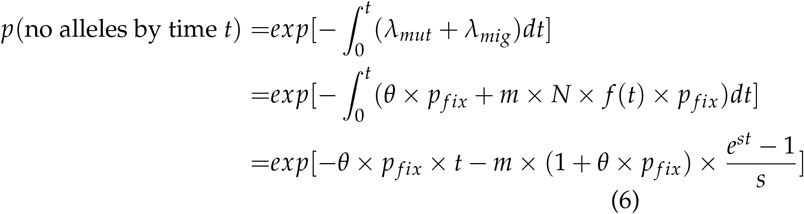

Then, the probability of the first allele entering via mutation is

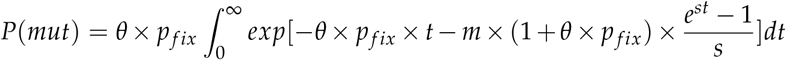

This roughly represents the probability of a locally-derived sweep. This approximation works well as *p_fix_* approaches 1 (i.e., *s* large) and if *θ* is not too high.

**Table S1.** Medians and 95% posteriors for parameter estimation in intra-macaque Simian-HIV populations. *M* is a composite estimate taken from multiplying the *m* distribution by the point estimate of predicted population size derived from the estimated *θ* (*M* = *m* × *θ/μ*). The parameter *c* represents the threshold for including mutations in haplotypes. *c* = 1 is the most lenient filter and *c* = 5 is the most strict filter. The results for *c* = 2 are also plotted in Table 1. See the Materials and Methods for the full description of *c*.

| | $c$ | $m$ | $M$ | $s$ | $\theta$ |
| --- | --- | --- | --- | --- | --- |
| Plasma v Gut | 1 | 0.00103 | 101 | 1.5 | 0.98 |
|  |  | (5e-05,0.291) | (4.9,28600) | (0.71,5.31) | (0.559,1.94) |
| Plasma v Lymph Node |  | 0.0267 | 2460 | 0.949 | 0.92 |
|  |  | (0.00285,0.448) | (262,41200) | (0.604,5.01) | (0.51,2.14) |
| Plasma v Gut | 2 | 0.000528 | 15.5 | 3 | 0.293 |
|  |  | (2.06e-05,0.0326) | (0.602,953) | (0.987,27.7) | (0.122,0.744) |
| Plasma v Lymph Node |  | 0.0271 | 573 | 1.24 | 0.212 |
|  |  | (0.00109,0.444) | (23,9400) | (0.783,3.19) | (0.11,0.44) |
| Plasma v Gut | 5 | 0.000537 | 8.05 | 1.75 | 0.15 |
|  |  | (1.84e-05,0.0145) | (0.276,218) | (1,15.8) | (0.103,0.296) |
| Plasma v Lymph Node |  | 0.0413 | 574 | 1.52 | 0.139 |
|  |  | (0.00103,0.447) | (14.4,6220) | (0.849,14.9) | (0.101,0.29) |

**Figure S1.**
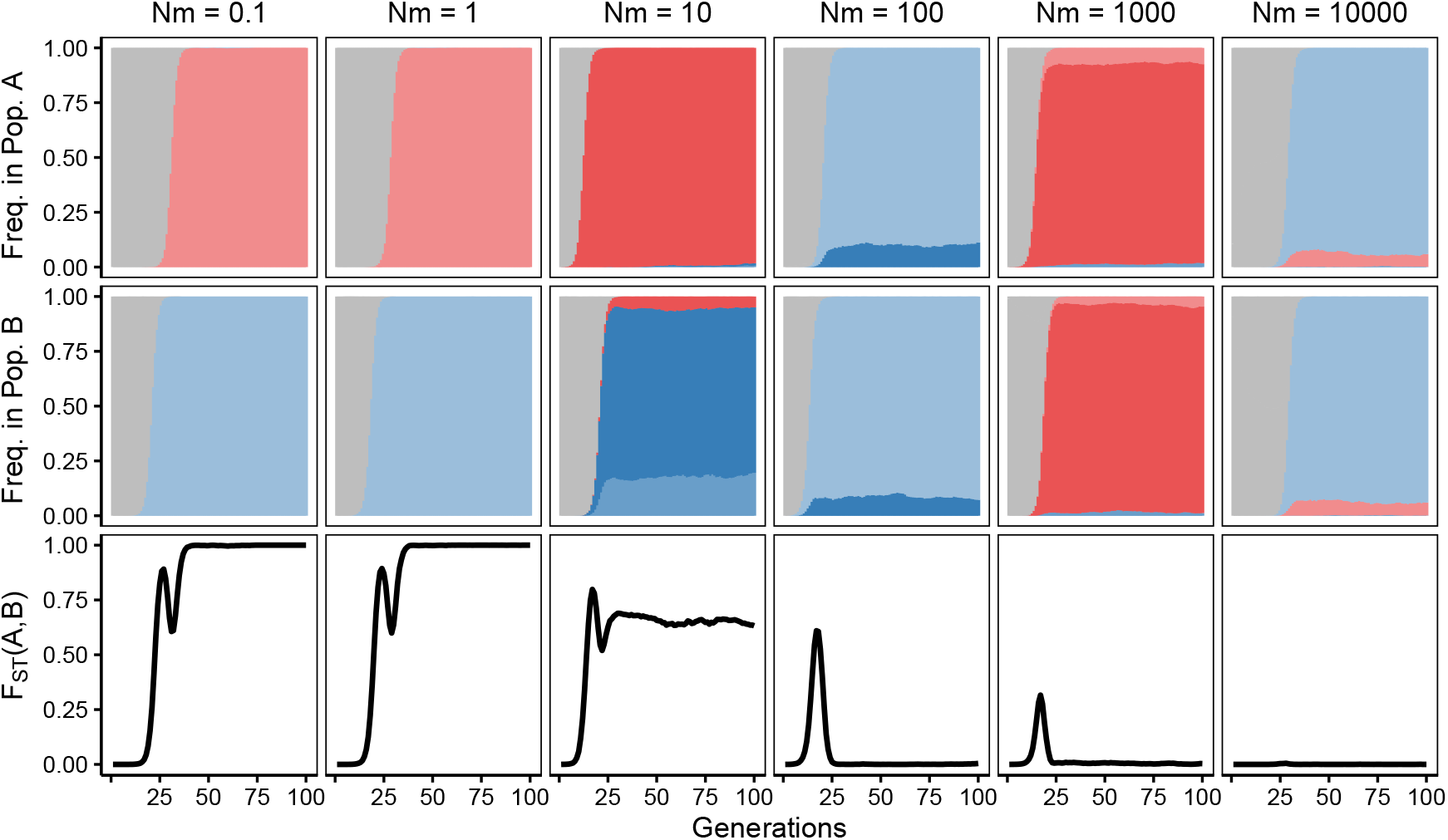
Patterns of neutral diversity over time. This figure is the analog to Figure 3 with lower population mutation rate (*θ* = 0.5). Correspondingly, the sweeps show characteristics of migrant-derived (rather than locally-derived) sweep patterns.

**Figure S2.**
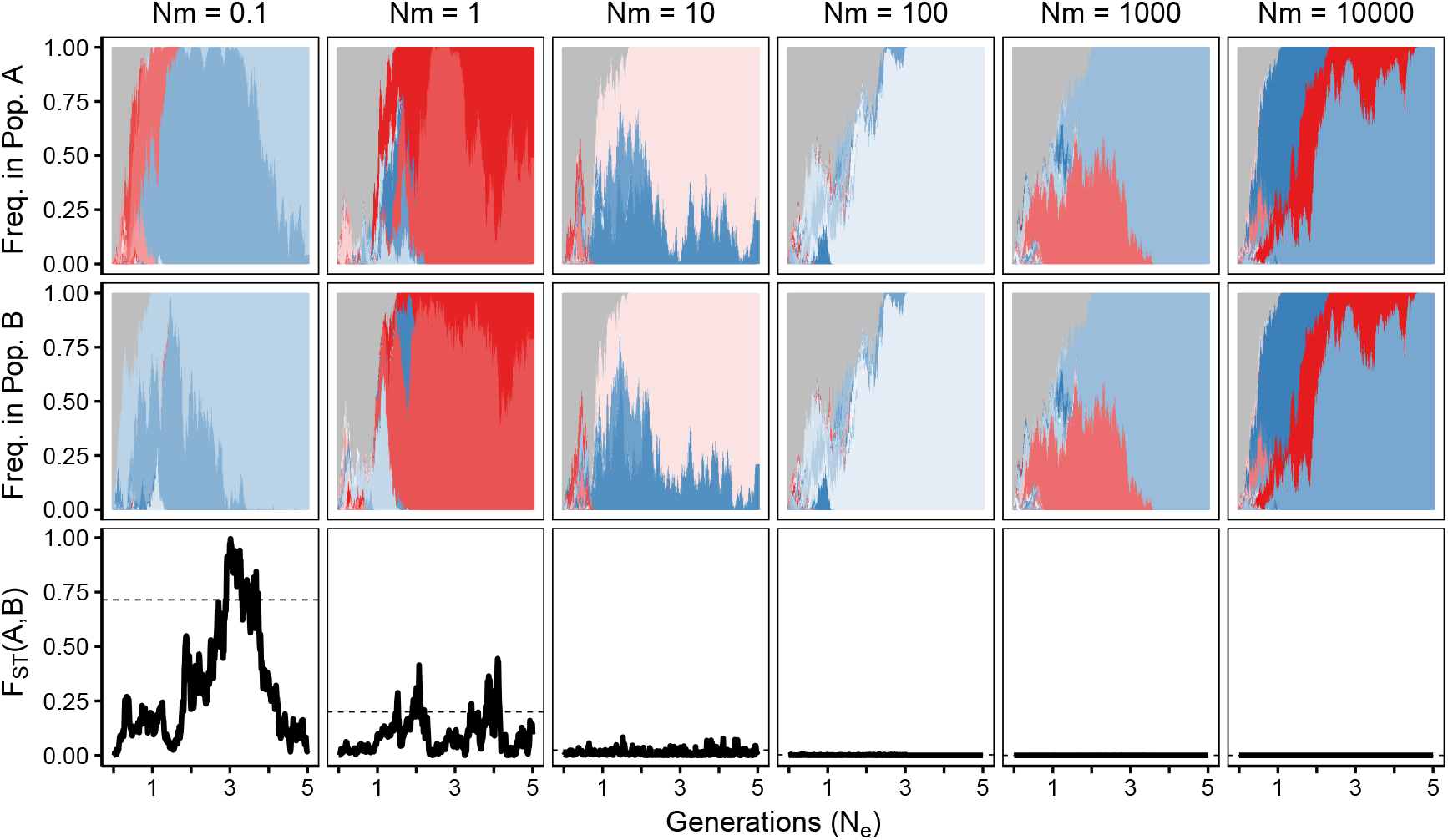
Patterns of neutral diversity over time. This figure is the analog to Figure 3 under neutrality (i.e. selection strength *s* = 0). Correspondingly, the timescale for alleles to reach high frequencies is now in units of 10^5^ (i.e., *N_e_*) generations instead of single generations. The dashed lines mark the result *F_ST_* = 1/ (4*Nm* + 1), and the figure caption is otherwise shared with Figure 3.

**Figure S3.**
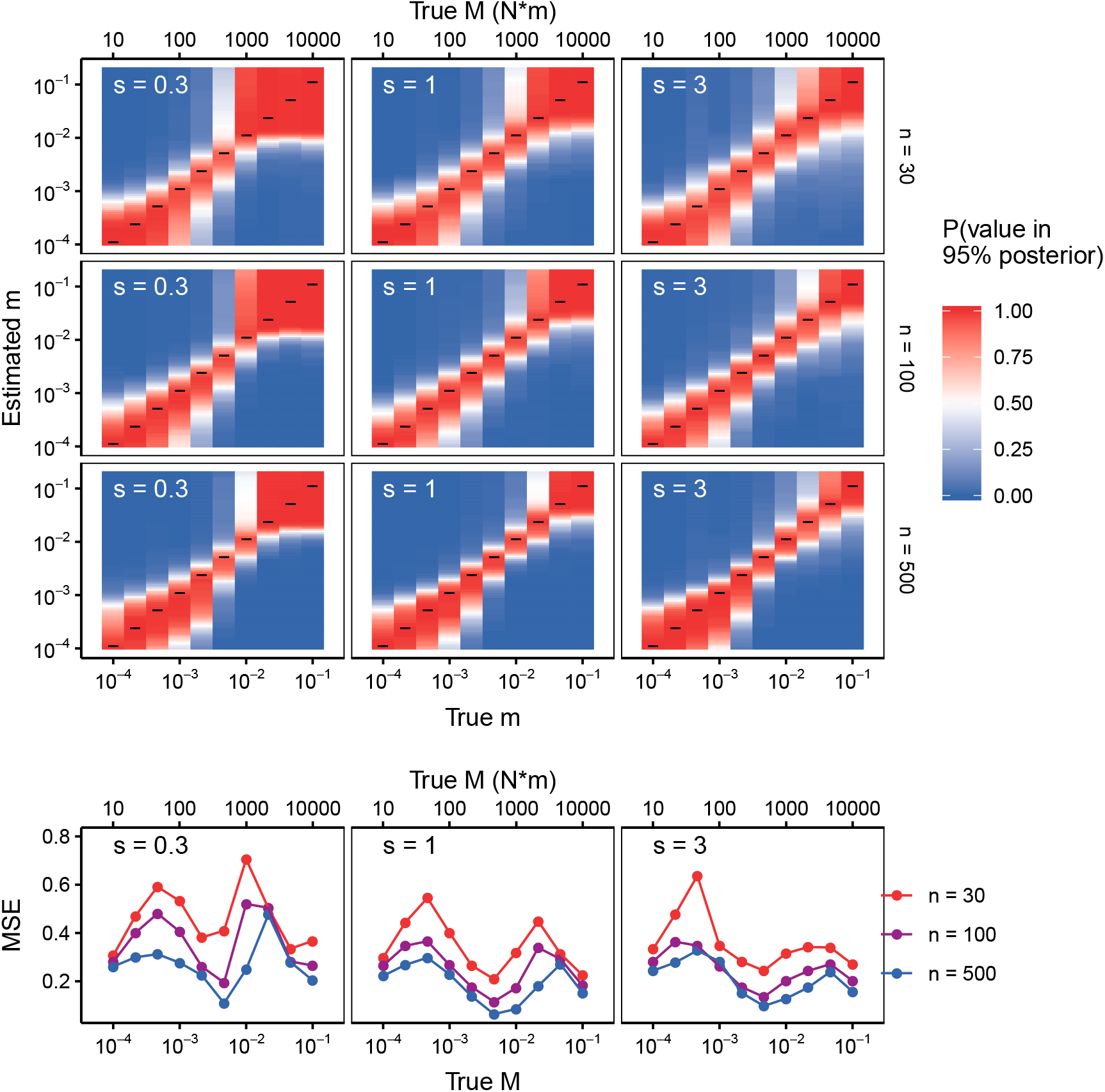
Increasing the sampling depth *n* decreases the average distance between the posterior and the truth. Similar to Figure S4, this figure displays the ability of the ABC procedure to recapture a test ‘Estimated m’ under different values of *s* and *M*, with different sampling depths per time point (*n*), as shown in rows. Increasing *n* results in estimation more focused around the *x* = *y* line. This is displayed in the bottom row as decreased log-MSE among the highest *n* simulations, across values of *s* and *m* (*N* = 10^5^, 200 replicates).

**Figure S4.**
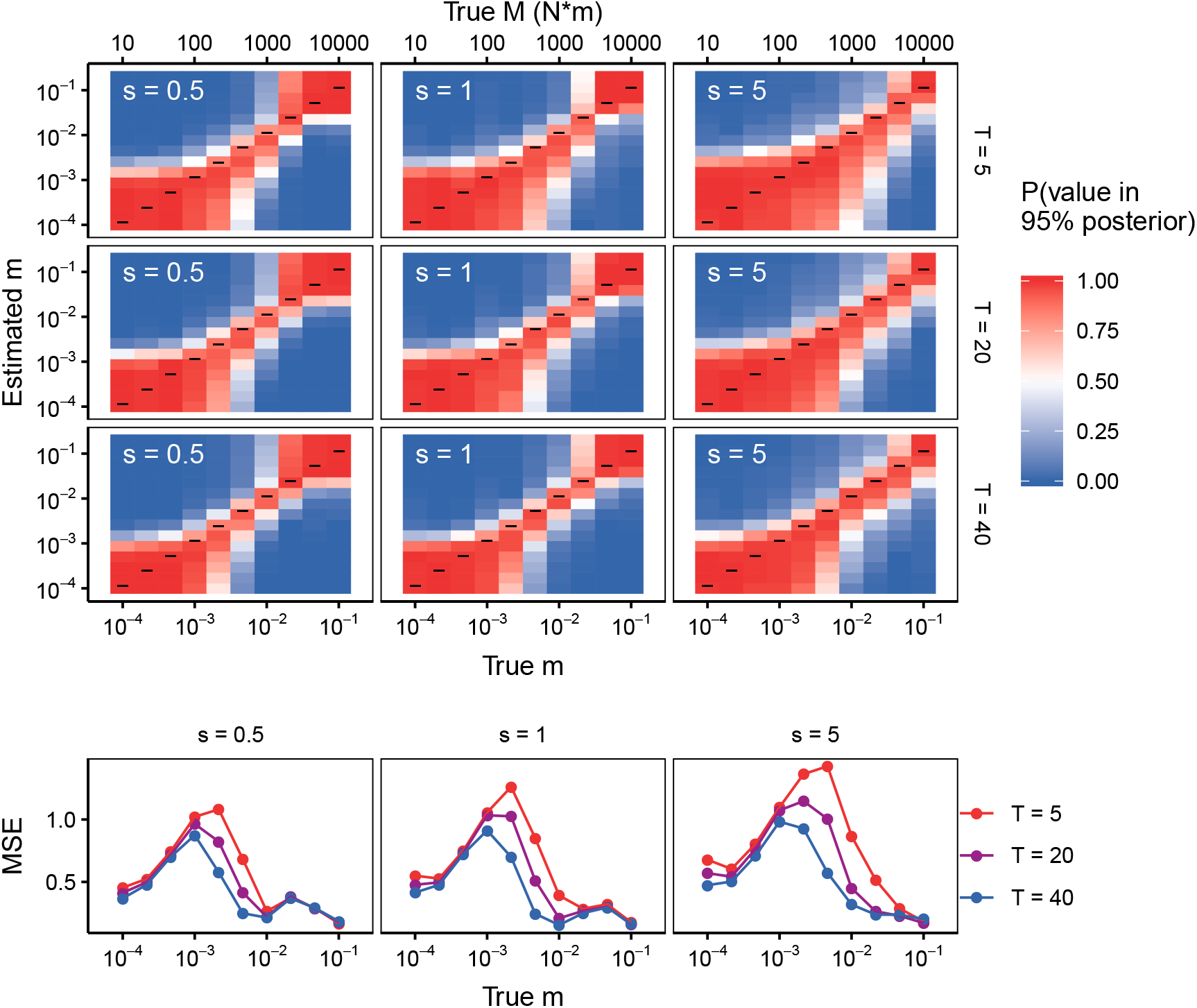
Increasing T decreases the average distance between the posterior and the truth. Similar to Figure, this figure displays the ability of the ABC procedure to recapture a test ‘Estimated m’ under different values of *s* and *M*, with different sampling time configurations, as shown in rows. Sampling time points were taken at generations 5, fixation, and fixation + *T*, with *T* = (5,20,40) Increasing *T* can result in estimation more focused around the *x* = *y* line, indicating precise and accurate estimation. This performance is displayed in the bottom row as decreased log-MSE among the highest *T* simulations, across values of *s* and *m*. This effect is most important for particular values of *T* (*N* = 10^5^).

**Figure S5.**
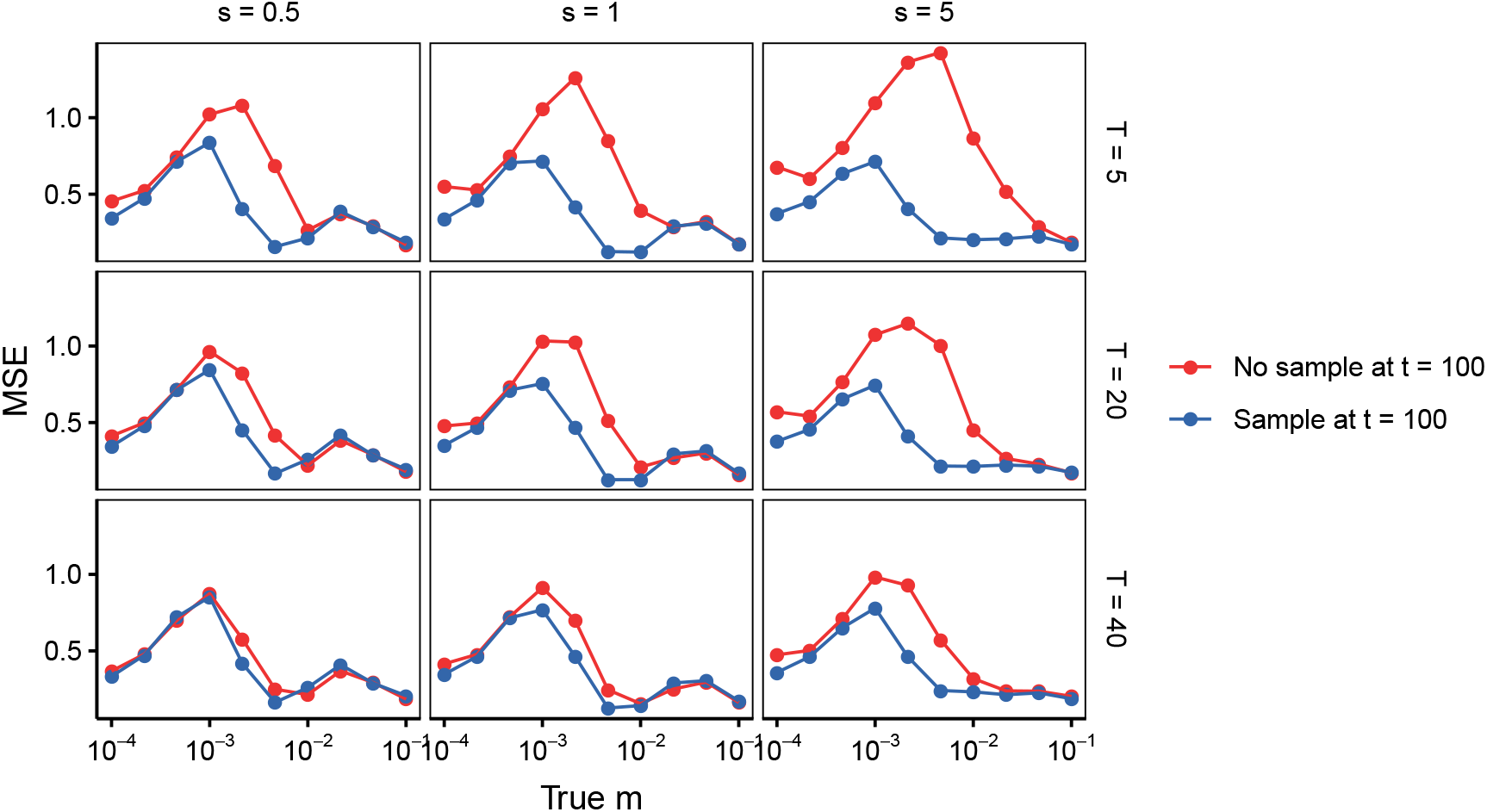
Adding one sample time point at generation 100 substantially improves fit, but significantly more when T is small. For different values of *T, s* and *M*, adding an additional time point at generation 100 decreases the log-MSE of the population. When *T* = 40, there is relatively little improvement, because the third sampling time point is likely already near *t* = 100 (*N* = 10^5^).

**Figure S6.**
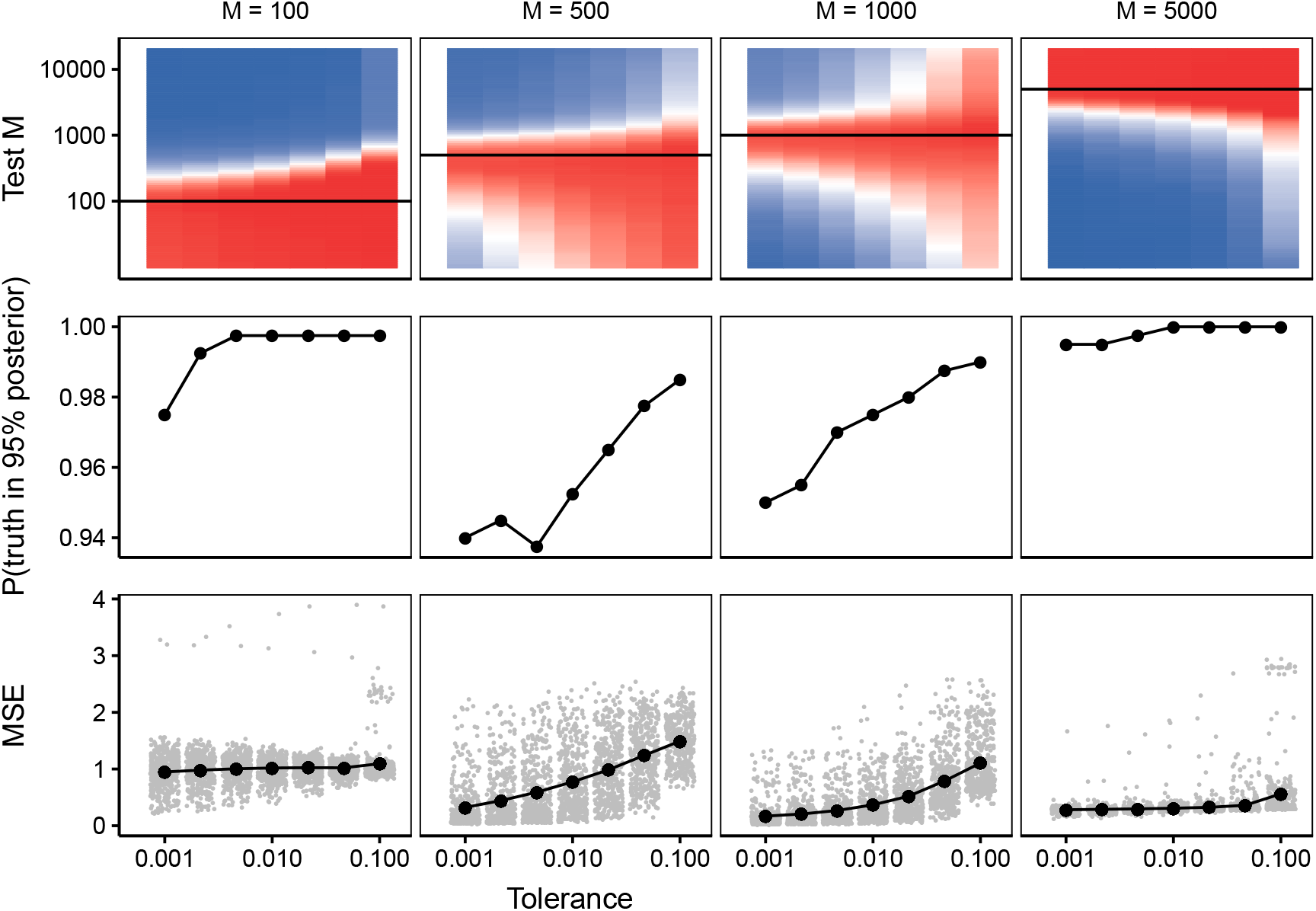
Decreasing the tolerance decreases the probability of finding the actual truth, but decreases the average distance between the posterior and the truth. The performance of the ABC procedure is evaluated in three different ways. In the top row, we show the probability of including different values of *M* within posteriors computed with different tolerances. The true value of *M* is shown with the black line. The blue to red probability spectrum indicates that lower tolerances include fewer false values of *M* within their corresponding 95% posteriors. In the second row, the proportion of trials with a given tolerance that successfully capture the true *M* decreases with the tolerance. In the third row, the log-MSE decreases as the tolerance is decreased. Each grey dot shows an individual simulation with a given *M* and the black dots display the median (*N* = 10^5^, *s* = 1).

**Figure S7.**
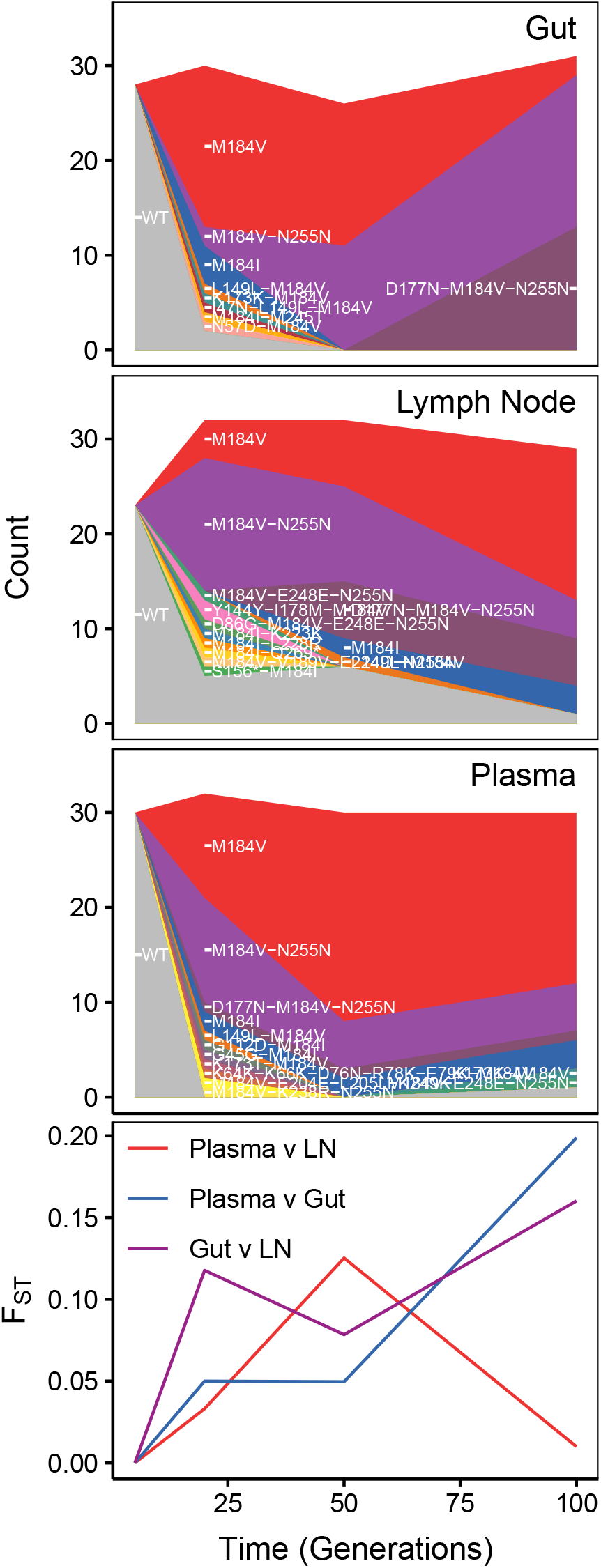
Dynamics of drug resistance fixation across space and time in a treated Simian-HIV population with lenient haplotype definitions. (*c* = 1). The top row shows diagrams of drug resistant haplotypes spreading in different sampling locations over time sampled at generations 7, 21,49 and 98 after the onset of selection via the drug FTC in the gut, lymph node and blood plasma. Mutations are included if they are observed in at least a single copy at or before a sweep is at least 70% complete (see Materials and Methods for full description of how haplotypes were defined)Each color represents a distinct lineage separated by at least one mutation. The bottommost panel plots pairwise *F_ST_* between pairs of sampling locations.

**Figure S8.**
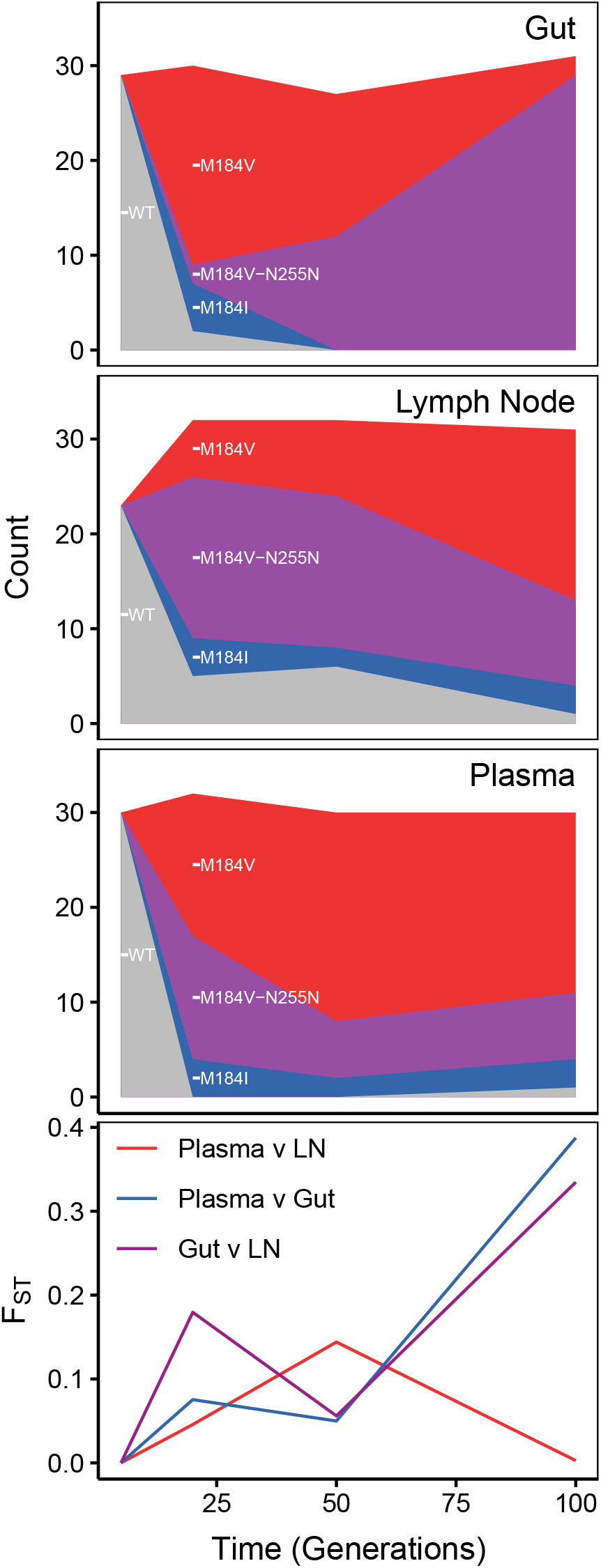
Dynamics of drug resistance fixation across space and time in a treated Simian-HIV population with strict haplotype definitions. (*c* = 5). The top row shows diagrams of drug resistant haplotypes spreading in different sampling locations over time sampled at generations 7, 21, 49 and 98 after the onset of selection via the drug FTC in the gut, lymph node and blood plasma. Each color represents a distinct lineage separated by at least one mutation. Mutations are included if they are observed in at least five copies at or before a sweep is at least 70% complete (see Materials and Methods for full description of how haplotypes were defined). The bottommost panel plots pairwise *F_ST_* between pairs of sampling locations.

**Figure S9.**
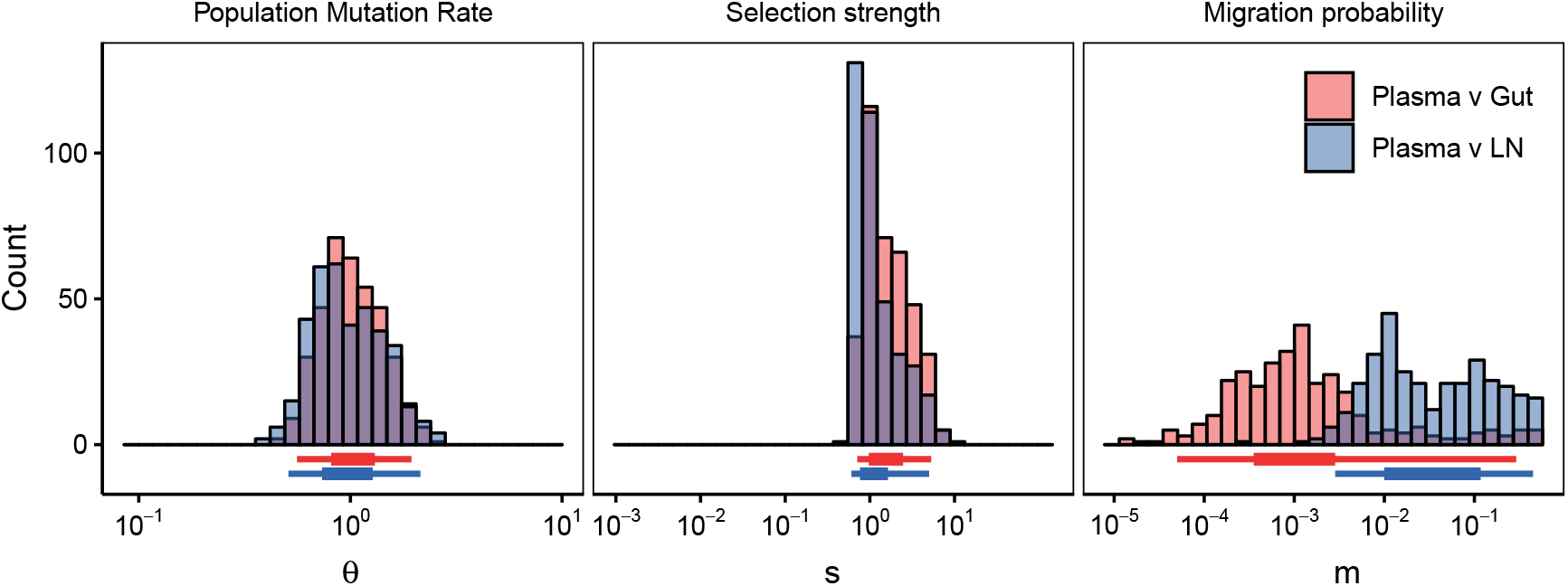
Estimation of population parameter rates from intra-patient Simian-HIV data sampled from different subpopulations with lenient haplotype definitions. (*c* = 1). The top row shows diagrams of drug resistant haplotypes spreading in different subpopulations over time sampled at generations 7, 21, 49 and 98 in the gut, lymph node and blood plasma. Each color represents a distinct lineage separated by at least one mutation. The resulting posteriors are given for the ABC procedure for comparisons between the plasma and gut (red) and plasma and lymph node (blue). 95% posteriors for these distributions are shown in Table S1.

**Figure S10.**
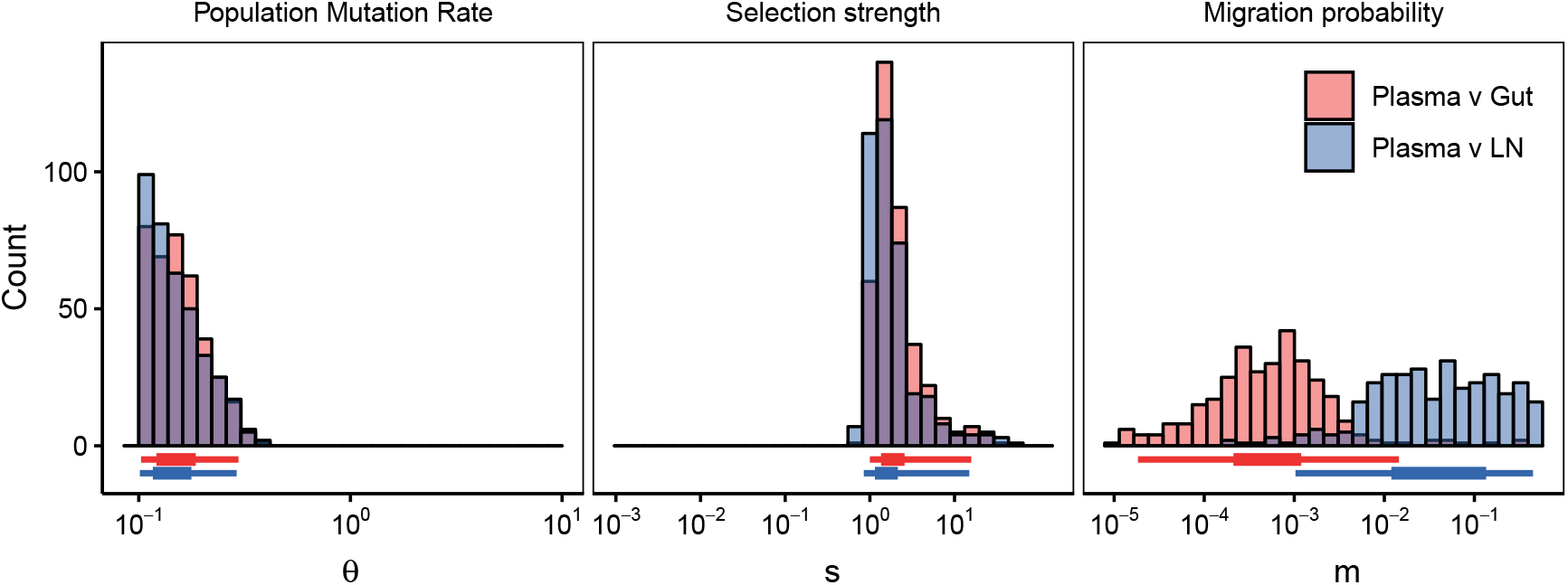
Estimation of population parameter rates from intra-patient Simian-HIV data sampled from different subpopulations with strict haplotype definitions. (*c* = 5). The top row shows diagrams of drug resistant haplotypes spreading in different subpopulations over time sampled at generations 7, 21, 49 and 98 in the gut, lymph node and blood plasma. Each color represents a distinct lineage separated by at least one mutation. The resulting posteriors are given for the ABC procedure for comparisons between the plasma and gut (red) and plasma and lymph node (blue). 95% posteriors for these distributions are shown in Table S1.

**Figure S11.**
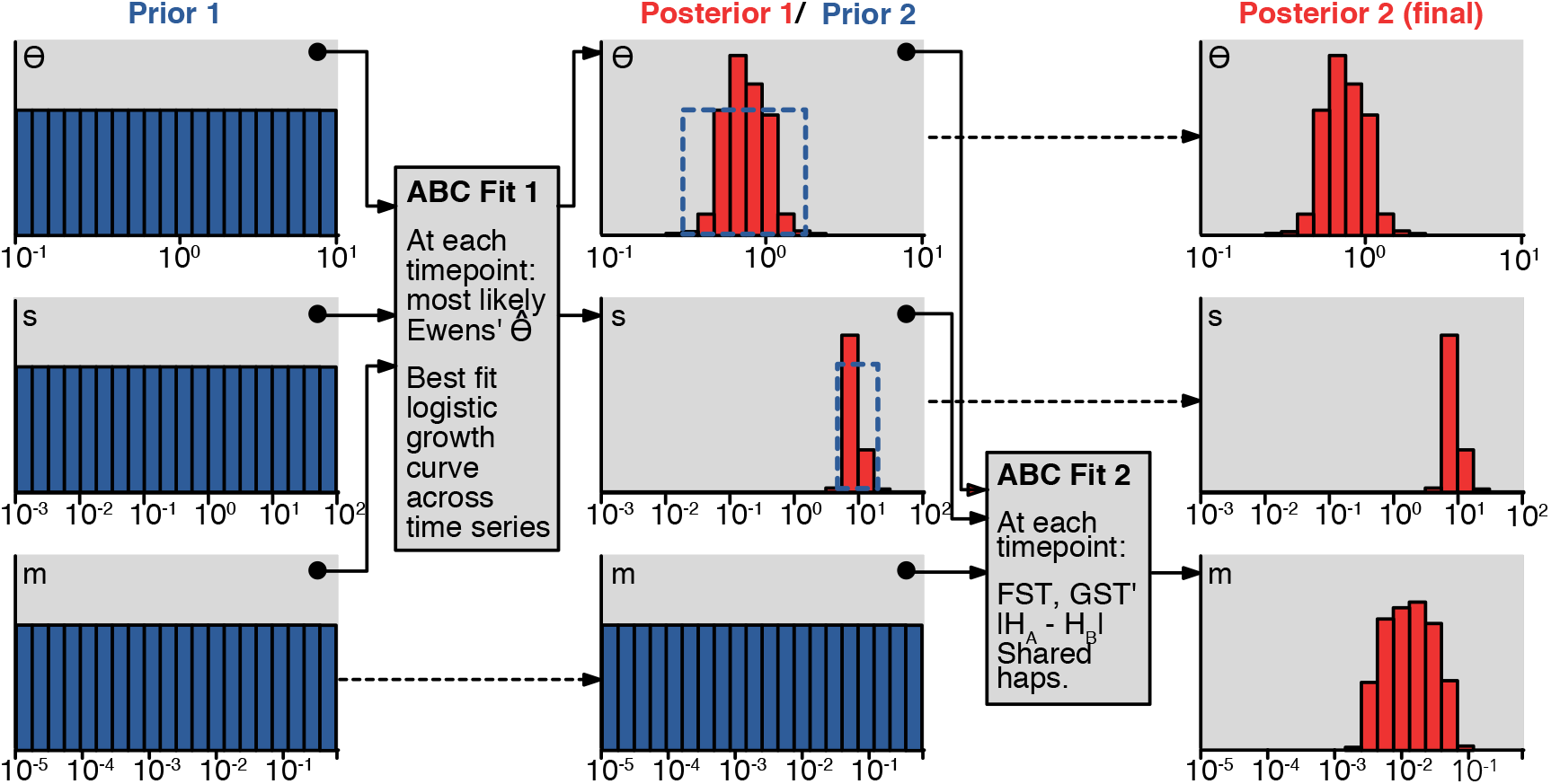
Schematic of 2-step approximate Bayesian computation (ABC) procedure. Priors for all three parameters are initially *log*_10_ uniform with *θ* ∈ (10^−1^,10^1^), *s* ∈ (10^−3^,10^2^), *m* ∈ (10^−5^,.5), and these priors are used to estimate posteriors over *θ* and *s* with summary statistics tailored for mutation and selection as shown in “ABC Fit 1.” The posteriors from the first ABC fit are used as priors for the second round ABC, while the posterior for *m* remains log_10_ uniform with *m* ∈ (10^−5^,.5). The summary statistics in the second fit are tailored for estimating migration. The final posteriors are *m* posterior from ABC 2 and the *s* and *θ* posteriors from ABC 1. For more details, see Materials and Methods, Approximate Bayesian Computation.

